# CAR T cell therapy selectively depletes disease-driving mutant calreticulin cells in xenotransplants and human organoid models of myelofibrosis

**DOI:** 10.64898/2026.01.28.702220

**Authors:** Alexandros Rampotas, Zoë C. Wong, Isaac Gannon, Charlotte Brierley, Yuqi Shen, Camelia Benlabiod, Ashlyn Chee, Saif Khan, Nawshad Hayder, Gordon Weng-Kit Cheung, Marina Mitsikakou, Eleanor Murphy, Mathieu Ferrari, Anna Bulek, Antonio Rodriguez-Romera, Lauren Murphy, Aude-Anais Olijnik, Manuel Rodriguez-Justo, Caroline Marty, Ian Hitchcock, Daniel Royston, Adam J. Mead, Abdullah O. Khan, Jonathan Lambert, Claire Roddie, Bethan Psaila, Martin A. Pule

## Abstract

Targeted immunotherapies have revolutionized outcomes for lymphoid malignancies, but success in myeloid neoplasms is limited by the lack of amenable targets and immunologically hostile tumor microenvironment (TME). Myeloproliferative neoplasms are chronic myeloid blood cancers, a third of which are driven by mutations in calreticulin (mutCALR). This yields a common neoepitope that binds to, and activates, the thrombopoietin receptor and results in display of the oncoprotein on the extracellular membrane of disease-driving cells, exposing a therapeutic vulnerability. Here, we present a first-in-class chimeric antigen receptor (CAR) T-cell therapy that specifically targets mutCALR+ cells, both *in vitro* and *in vivo.* The CAR T-cell therapy selectively depleted mutCALR+ stem cells from patients with myelofibrosis while sparing healthy stem cells, and improved survival in mutCALR leukemia xenografts. To mimic myelofibrotic marrow, we developed a bespoke human ‘chimeroid’ model and showed no decrease in the potency of CAR T cell-mediated target cell killing even in a fibrotic tumor microenvironment. We also devised a method to boost cell surface expression of mutCALR in CD34+ cells isolated from patients with accelerated/blast phase MPN (defined as >10 % blasts in peripheral blood or bone marrow), enhancing CAR T targeting. This study presents a therapeutic with potential to eradicate mutCALR-driven malignancies and highlights an innovative strategy to evaluate blood cancer-targeting immunotherapies in a relevant TME.

**One Sentence Summary:** A first-in-class CAR T-cell therapy targeting mutant calreticulin selectively depletes malignant stem cells *in vivo* and in fibrotic human organoids.

## INTRODUCTION

Targeted cancer therapies were first pioneered in BCR-ABL positive chronic myeloid leukemia, transforming a rapidly fatal diagnosis to a condition where most patients now achieve long-term remission or cure (*1*). Similarly, targeted immunotherapies have dramatically improved outcomes for patients with lymphoid malignancies (*2*)(*3*). Meanwhile, success in developing a cancer-selective therapy for BCR-ABL negative myeloproliferative neoplasms (MPNs) has remained elusive, and these cancers remain largely incurable. MPNs are a group of chronic myeloid blood cancers that arise following the acquisition of a gene mutation in a hematopoietic stem cell (HSC). In around a quarter of patients, the driver mutation affects the gene encoding the endoplasmic reticulum (ER) chaperone protein calreticulin (mutCALR) (*4*)(*5*). Over time, mutCALR-driven MPNs can progress to myelofibrosis (MF) (*6*), a severe malignancy where inflammation and fibrosis cause bone marrow failure, resulting in cytopenias and a markedly reduced life expectancy of 5-7 years. Up to 1 in 5 patients with MF transform further to accelerated/blast phase MPN (AP/BP-MPN), curtailing life expectancy to 6-12 months (*7*).

Current MPN therapies largely comprise cytoreductive agents that non-selectively reduce myeloproliferation and symptoms but do not alter the dynamics of progression (*8*). Therefore, there remains an urgent need for cancer cell-selective targeted therapies to improve outcomes for patients with higher risk disease. While several different *CALR* mutations occur, all known pathogenic mutations create a +1 base-pair frameshift in exon 9, generating a truncated CALR protein with a novel C-terminus. This leads to two critical consequences: Firstly, formation of an aberrant complex with the thrombopoietin receptor (TpoR) causing oncogenic cytokine receptor signaling, and secondly, loss of the ER-retention signal enables the mutCALR-TpoR complex to migrate to the cell membrane of TpoR-expressing cells – which includes cancer-initiating stem cells and fibrosis-driving megakaryocytes (*9*)(*10*). This presents a therapeutic opportunity to selectively target the key cellular mediators of pathobiology in mutCALR+ cancers, allowing normal hematopoiesis to recover.

Exploiting the opportunity offered by the cell surface-expressed cancer neoantigen, a monoclonal antibody (*11*) and a T-cell engaging (TCE) bi-specific antibody (*12*) recently entered phase I studies. However, myelofibrotic marrow represents a challenging environment for immunotherapy due to the dense extracellular matrix and immunologically hostile cellular and cytokine milieu (*13*). Chimeric antigen receptor T (CAR T) cell therapies have demonstrated lasting remissions in patients with refractory B-cell (*3*) and plasma cell malignancies (*14*), and can be more potent than other targeted immunotherapy approaches (*15*). Their ability to actively penetrate diseased tissue may be particularly useful in overcoming the fibrotic and immunosuppressive microenvironment marrow associated with mutCALR MPNs.

Here, we present preclinical characterization of a CAR T cell therapeutic that recognizes the mutant-specific C-terminus of mutCALR. We show selective depletion of mutCALR+ cells *in vitro*, in a bespoke human bone marrow organoid model and *in vivo*, including against samples from patients with a range of CALR mutations and additional co-mutations. We demonstrate that the CAR T cells reduce the mutant clone burden in healthy and fibrotic marrow niches, and we also present a novel strategy to boost mutCALR expression and enhance killing in low target-expressing cancer cells. These findings set the stage for a new cancer-specific therapy for patients with a high risk and currently incurable blood malignancy and validate a broadly relevant experimental platform for evaluating blood cancer-targeting immunotherapies in a relevant human tissue environment.

## RESULTS

### Characterization of a mutCALR-specific single-chain variable fragment (scFv)

We designed a single-chain variable fragment (scFv) specific to the mutant C-terminus of the protein that is common to all known pathogenic mutations (RPG4-Fc, Fig. 1A). Indirect ELISA and surface plasmon resonance confirmed selectivity for the mutant protein (Fig. 1B & C) and optimal binding stability (Fig. S1). To assess selectivity in a cellular context, we leveraged the requirement for co-expression of TpoR to enable cell surface presentation of mutCALR. A panel of human (Marimo, UT-7-TPO) and murine (Ba/F3) cell lines was engineered to express either wild-type or mutCALR, with or without the human TpoR (Fig. 1D). RPG4-Fc only bound murine Ba/F3 cells if both mutCALR and human TpoR were co-expressed (Fig. 1E). Evaluation of binding to disease-relevant, human MPN cell lines showed binding to Marimo cells, a human mutCALR+ myeloid leukemia cell line, and this was increased following TpoR over-expression (Fig. 1E). Similarly, RPG4-Fc only bound the TpoR-expressing megakaryoblastic (wild-type *CALR*) leukemia UT-7 cell line after transduction with mutCALR, and binding was further enhanced with TpoR over-expression (Fig. 1E).

**Fig. 1.**
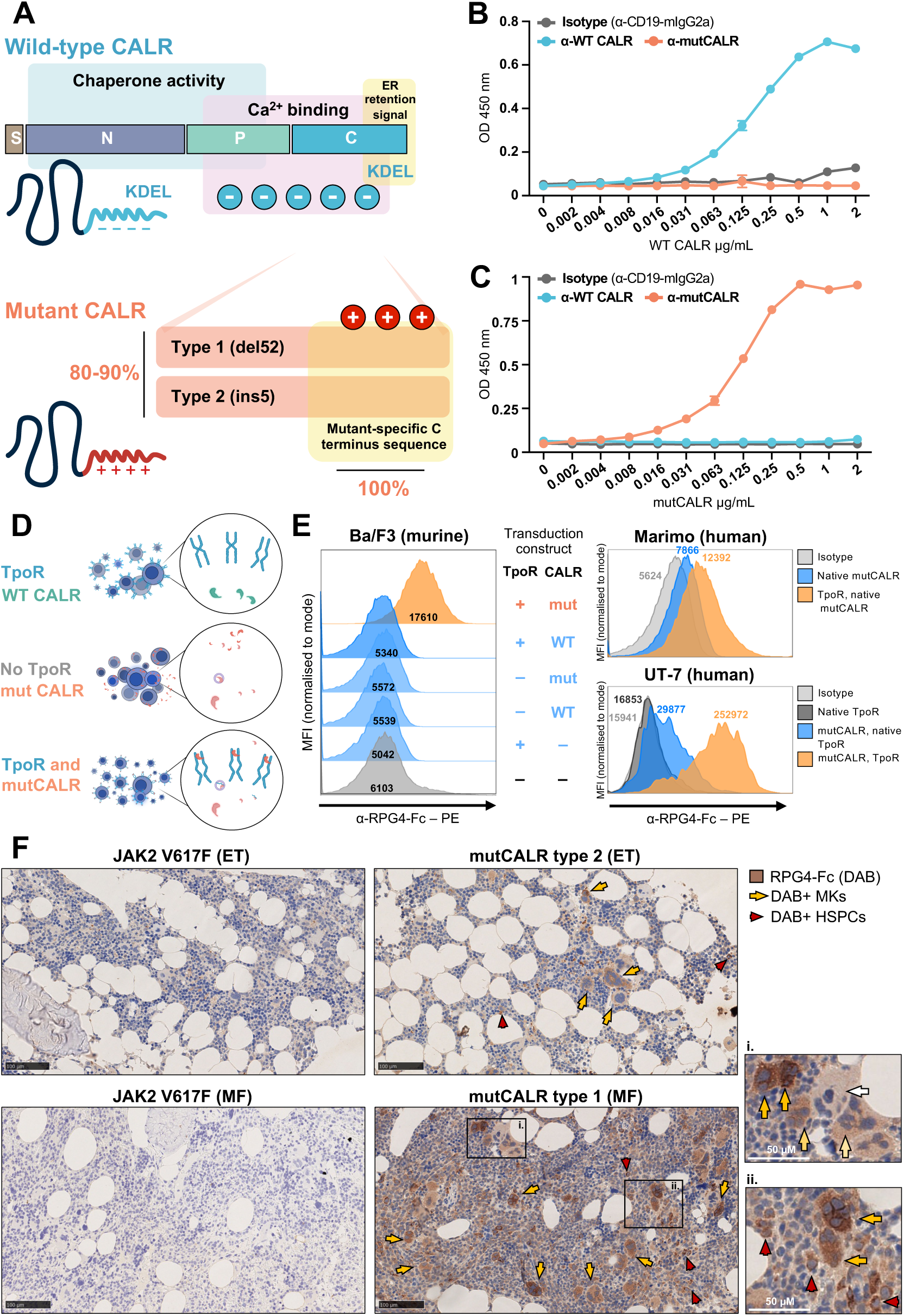
Development and validation of RPG4 single chain variable fragment (scFv) against mutant calreticulin (mutCALR). **(A)** Graphical representation of how all known oncogenic mutations in *CALR* result in the same common mutant C-terminus of the protein, with an altered electrostatic charge that causes formation of a complex with the thrombopoietin receptor. Loss of the KDEL endoplasmic reticulum (ER) retention signal allows the complex to exit the ER and be expressed on the cell surface. 80-90% of patients have either a Type 1 (52 base pair deletion) or Type 2 (5 base pair insertion) mutation, which both result in a frameshift mutation, while other mutations are classified as type-1 or type-2 like. **(B, C)** Indirect ELISA for RPG4 (1 μg/μL) using various concentrations of purified wild-type calreticulin (WT CALR, **B**) or the mutant C terminus of mutCALR (**C**). Antibodies against WT CALR and against human CD19 were used as controls (n=3). **(D)** Schematic showing how mutCALR is only presented on the cell surface of cells that express the thrombopoietin receptor (TpoR). **(E)** Flow cytometry analysis of engineered murine Ba/F3 cells and human Marimo and UT-7 cells stained with RPG4-Fc and a secondary anti-murine IgG2a PE-conjugated antibody. Analysis of non-transduced Ba/F3 cells, as well as panel of cells expressing wild-type or mutCALR with/without co-transduction of the TpoR as indicated. **(F)** Immunohistochemistry staining using the RPG4 scFv of bone marrow biopsies from patients with either essential thrombocythemia (ET) or myelofibrosis (MF) driven by Type 1 or Type 2 *CALR* mutations, or JAK2V617F as a negative control. Brown colour (3, 3’-diaminobenzidine, DAB) indicates positive staining. Arrows indicate DAB+ megakaryocytes (yellow) and HSPCs (red). **(i)** Yellow to white arrows reflect intensity (strong to weak/negative) of RPG4-Fc staining on megakaryocytes. **(ii)** Red arrows highlight small mononuclear cells, consistent with HSPCs.

Specificity in human bone marrow was confirmed by immunohistochemical staining of bone marrow biopsies from patients with mutCALR+ MPNs, or JAK2V617F+ MPNs as a negative control (n=8). RPG4-Fc showed positive staining of the TpoR-expressing cell types (megakaryocytes and hematopoietic stem/progenitor cells [HSPCs]) in bone marrow of mutCALR+ but not JAK2V617F+ biopsies (Fig. 1F, fig. 1e & 1f, Suppl. Table S1). Equivalent staining was seen using RPG-4-Fc to that using the commercially available anti-mutant CALR ‘CAL2’ immunohistochemistry antibody (*16*), with no difference in staining of biopsies from patients with Type 1 and Type 2 variants (fig. S1e). mutCALR positivity was observed on both mature megakaryocytes and CD34+ HSPCs (fig. S1f).

### RPG4 CAR T cells exhibit potent and specific killing of mutCALR/TpoR+ target cells

RPG4-Fc was incorporated into a 41BB-ζ CAR construct (*17*) (RPG4-BBζ) with the sort-suicide gene RQR8 (*18*) in a bicistronic vector (Fig. 2A). RPG4 CAR T cells were tested against dilutions of plate-bound, purified mutCALR protein C-terminus with CD19 CAR T cells as a negative control. Dose-dependent upregulation of the activation marker CD25 was observed in response to mutCALR, with no activation of CD19 CAR T cells (Fig. 2B). At an effector:target (E:T) ratio of 1:1 and with read-outs performed at 72 hours (hrs), RPG4 CAR T cells killed Ba/F3 cells and released IFNγ and IL2 only when exposed to Ba/F3 cells that co-expressed mutCALR and TpoR, but not TpoR alone or wild-type CALR (Figs. 2C and D). Similarly, RPG4 CAR T cells killed and released IFNγ/IL2 in response to mutCALR-positive Marimo and UT-7 cells, with enhanced killing with TpoR over-expression (Fig. 2E, F, fig. S2). At a lower E:T ratio of 1:4, killing was only marginally decreased at 72 hrs and was similar to the 1:1 ratio after 6 days (Fig. S3). Upregulation of early and late activation markers (CD69 and CD25) and T-cell expansion were observed in response to mutCALR+ target cells, mirroring killing and cytokine release (fig. S4a-d).

**Fig. 2.**
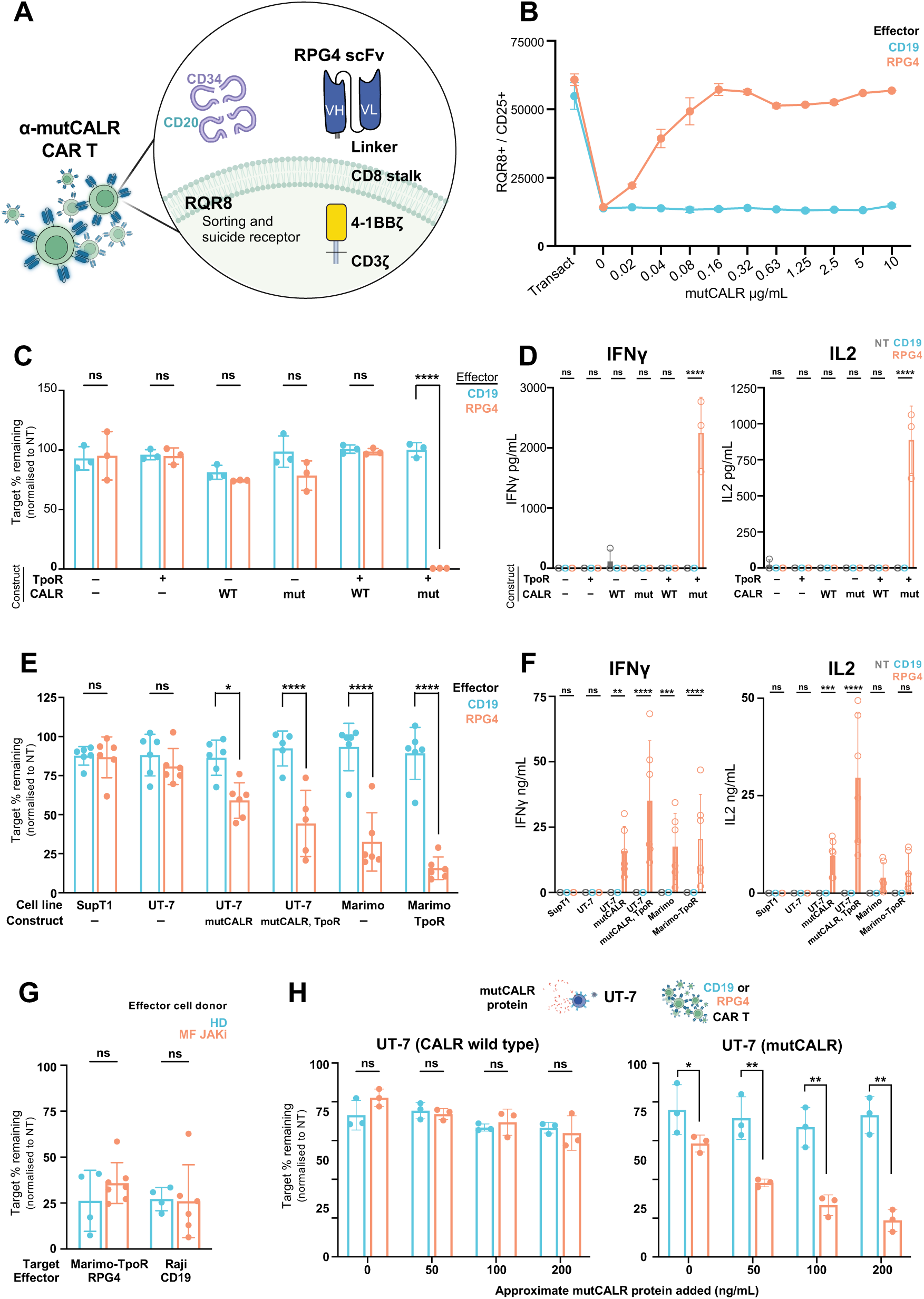
Development and validation of mutCALR-targeting CAR T cell therapy. **(A)** Graphical representation of CAR construct incorporating RPG4 binder. CAR construct used is a 2nd generation CAR with 4-1BB co-stimulatory domain and CD8 stalk as transmembrane and hinge domain. It also carries the RQR8 receptor which serves as a marker and ‘off-switch’. **(B)** Assessment of RPG4 CAR T sensitivity with a plate bound assay utilising increasing concentrations of the mutCALR C-terminus (n=3). **(C)** Flow cytometry-based killing assays using RPG4 CAR T against target positive and negative Ba/F3 cells in an effector:target ratio of 1:1. Ba/F3 cell numbers were quantified at 72 hours (hrs) and normalised to the condition where non-transduced (NT) T cells were added. CAR T cells were generated from healthy donors (n=3). Human CD19-targeting CAR T were used as a negative control. **(D)** Cytokine release measured by ELISA for IFNγ and IL2 24 hrs after addition of effector cells. **(E)** Killing assay and **(F)** IFNγ and IL2 secretion of non target-expressing (SupT1, UT-7-non-transduced, NT), low-expression (UT-7 mutCALR, Marimo NT) and high-expression (UT-7 mutCALR/TpoR, Marimo-TpoR) human cell lines using RPG4 CAR T generated from healthy donors (n=6 donors). (**G**) Comparative killing efficacy of CAR T cells generated from young healthy donors (n=4, 2 donors) and patients with myelofibrosis (MF) who were receiving JAK inhibitor (JAKi) therapy at the time of blood sampling (n=7 patients), tested at a 1:1 effector:target ratio. RPG4 and CD19 CAR T cells tested against Marimo-TpoR and CD19+ Raji cells, respectively. **(H)** Left - schematic showing experimental set up to test whether secreted mutCALR causes off-target cytotoxicity of non-CALR mutant cells via binding to TpoR. Right - target cells remaining 72 hrs after addition of RPG4 CAR T cells and CD19 CAR T cells (negative control) in conditions shown in the schematic (n=3). Comparisons made using 2-way ANOVA. * (p<0.05), ** (p<0.01), *** (p<0.001), ****(p<0.0001).

Chronic re-stimulation assays were performed to assess functionality under increased stress. RPG4 CAR T cells exhibited strong control of target cells after 3 rounds of co-culture, with continued, yet diminishing, control (fig. S4e) and a reduction in CAR T cell counts observed following round 5 (fig. S4f). Immunophenotyping after rounds 0, 1, 3 and 5 revealed an enrichment of naïve and T stem cell-like memory (Tscm) populations (CCR7+/CD45RA+/CD95+/CD62L+, fig. S4g). RPG4 CAR T cells remained non-exhausted until round 5, after which the majority of T cells expressed two or more exhaustion markers (PD-1, TIM-3, and LAG-3; fig. S4h). To assess transient acute re-stimulation, CAR T cells were exposed to target cells for 3 days followed by 4 days of rest in monoculture (fig. S5a). Continued control of target cells was observed with the CAR T cells from all 4 healthy donors, with ongoing CAR T cell expansion at assay completion (fig. S5b – S5d).

### CAR T cells generated from myelofibrosis patients on JAK inhibitor therapy have no impairment in function

T cells from patients with MPNs have increased expression of inhibitory receptors (*19*), and patients with myelofibrosis are frequently treated with JAK inhibitors which further inhibit T cell function (*20*). We therefore tested the function of CAR T cells generated from myelofibrosis patients receiving either ruxolitinib or momelotinib – two inhibitors of JAK1 and JAK2 that are licensed as first-line therapies for myelofibrosis. There were no significant differences in functionality of CAR T cells generated from these patients in comparison to CAR T cells derived from healthy donors (Fig. 2G).

### RPG4 CAR T cells do not target bystander TpoR-expressing, non-mutCALR cells

Secreted mutCALR could theoretically bind to TpoR-expressing wild-type cells and cause on-target, off-tumor cytotoxicity. It has been previously demonstrated that mutCALR only binds to TpoR expressed on mutCALR+ cells. (9). To test this, we investigated whether wildtype “bystander” TpoR-expressing cells could be bound by secreted/soluble mutCALR, and therefore inadvertently killed by RPG4 CAR T cells. Addition of CALRdel52-Fc supernatant (fig. S5f) to TpoR-expressing and wildtype UT7 cells did not induce CAR T cell-mediated killing and no CALRdel52 was detectable on the cell surface using RPG4-Fc as an antibody (fig. S5g). In contrast, mutCALR+ UT7 cells bound CALRdel52-Fc protein, and this binding led to enhanced RPG4 CAR T cell- killing in a dose-dependent manner (Fig. 2H). These findings are consistent with prior data showing preferential mutCALR binding to mutCALR+ TpoR-positive cells (*10*) and indicate that secreted mutCALR is unlikely to cause off-target killing of non-mutant TpoR-expressing cells.

### RPG4 CAR T selectively targets mutCALR+ myelofibrosis stem/progenitor cells

We next sought to test whether RPG4 CAR T cells could selectively deplete CALR mutant HSPCs from a cohort of patients with MPNs. RPG4 CAR T cells were co-cultured with HPSCs cells from patients with type 1 (52-base pair [bp] deletion), type 2 (5bp insertion), type 1-/2-like mutCALR+ myelofibrosis, or JAK2V617F-driven myelofibrosis as a negative control (n = 14, Suppl. Table S1). RPG4 CAR T cells eliminated between 60-75% of HSPCs from mutCALR myelofibrosis patients (Fig. 3A), including from patients with additional, high molecular risk mutations (Suppl. Table S1). No significant difference in killing was observed between type-1 or type-2 mutCALR subtypes (Fig. 3A), and no activation of CAR T cells or off-target cytotoxicity occurred in response to JAK2V617F+ myelofibrosis cells (Fig. 3A & 3B).

**Fig. 3.**
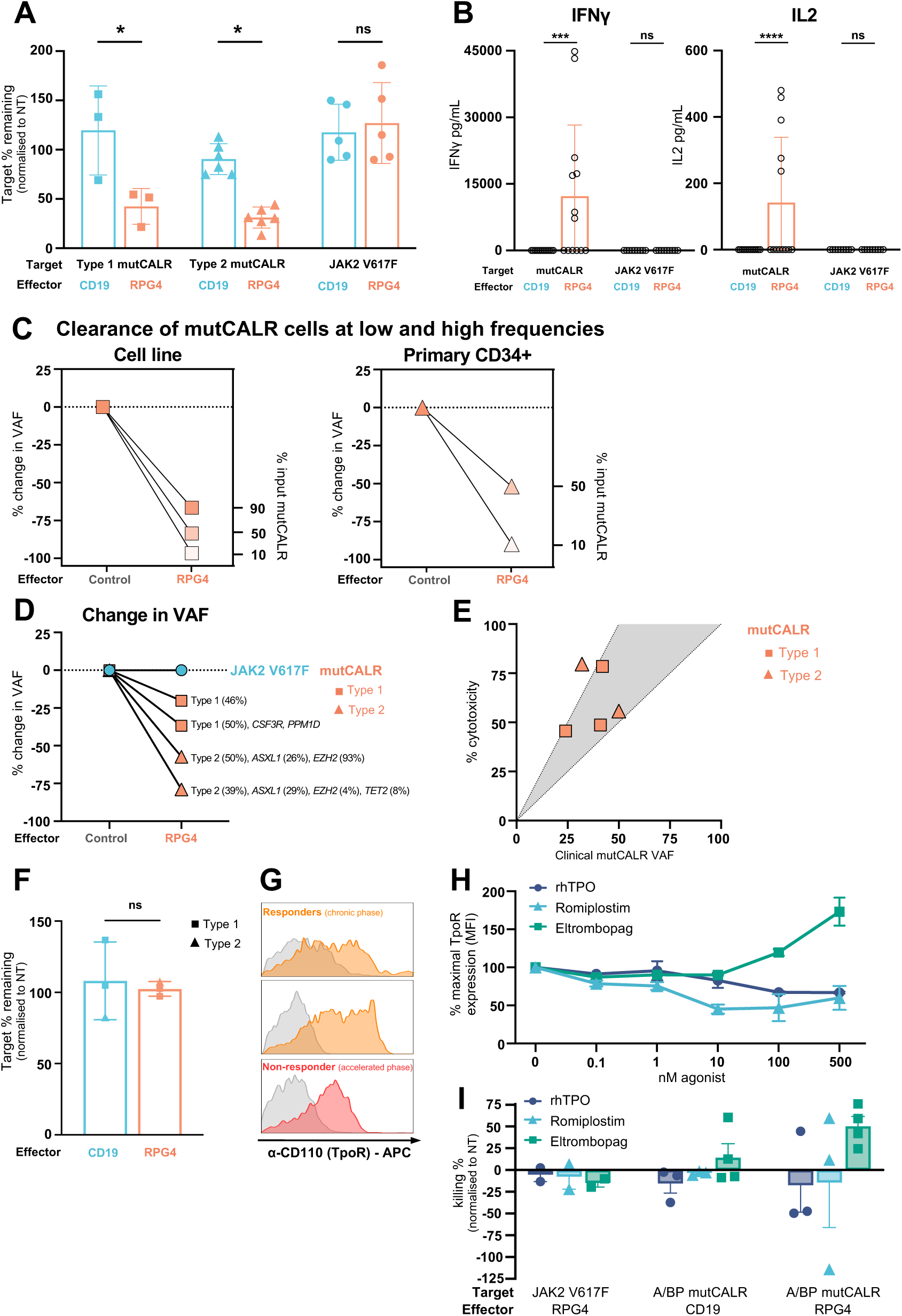
mutCALR-directed RPG4 CAR T cells deplete malignant CD34⁺ stem/progenitor cells from patients with myelofibrosis. **(A)** Target cell killing in co-cultures of RPG4 CAR T cells with primary CD34⁺ cells from patients harbouring mutCALR Type 1 (n=3 patients, squares) or Type 2 (n=6 patients, triangles) mutations. Primary CD34+ cells from patients with JAK2 V617F+ myelofibrosis were used as controls for off-target cytotoxicity (n=5 patients, circles). CD19 targeting CAR T cells (blue) were used as controls. **(B)** Cytokine secretion (IFNγ and IL2) measured in supernatant from CAR T cell co-cultures with primary cells as indicated in 3a. Samples were tested in duplicate or triplicate (n= 8, 2-3 biological replicates). (**C)** Change in variant allele frequency (VAF) following addition of CAR T cells to mixtures of mutant and wild-type cells to test killing efficacy at low (10%) and high (50%, 90%) target cell frequencies. Data for cell line (Marimo/-TpoR cells, left) and patient-derived CD34+ cells (Type 2, right) shown. VAF was measured by digital droplet PCR (ddPCR) and normalised to a control condition (non-transduced T cell or CD19 CAR T cell). **(D)** Change in mutCALR VAF following co-culture of cells from patients with mutCALR+ myelofibrosis, showing selective and potent depletion of mutCALR clones, including for patients with additional high molecular risk mutations (e.g. *EZH2*, *ASXL1*, *CSF3R*). JAK2V617F+ sample used as a control (showing no change in VAF) **(E)** Between 50 to >75% cytotoxicity was observed Type 1 and Type 2 mutCALR samples, correlating with clinical VAF. Shaded area on plot indicates 100% cytotoxicity for homozygous or heterozygous mutations. Zygosity of mutations for individual patients is unknown, and for heterozygous mutations a VAF of 25% would mean that 50% of cells bear mutCALR, therefore 50% cytotoxicity would reflect 100% killing of all target-positive cells. For homozygous mutations, a VAF of 25% would mean that 25% of cells bear mutCALR, therefore 25% cytotoxicity would reflect 100% killing of all target positive cells. **(F)** Accelerated/blast phase (AP/BP) CD34+ cells are not effectively targeted by RPG4 CAR T. **(G)** Thrombopoietin receptor (TpoR) expression for CD34+ cells from two chronic phase myelofibrosis samples vs. an accelerated phase MPN sample. Non-responders (red) have decreased TpoR expression at the cell surface in comparison to responders (orange). MFI is normalised to mode. **(H)** % maximal TpoR expression (mean fluorescence intensity, MFI) for Marimo-TpoR cells after stimulation with increasing concentrations of the TpoR agonists recombinant human TPO (rhTPO), romiplostim or eltrombopag. **(I)** Change in % killing of primary CD34+ cells isolated from patients with *JAK2* V617F+ myelofibrosis or mutCALR+ AP/BP MPN following overnight TpoR stimulation with rhTPO (purple), romiplostim (blue), or eltrombopag (green). Significance levels for 2-way ANOVA: p < 0.05 (*), < 0.01 (**), < 0.001 (***), <0.0001 (****).

Patients with myelofibrosis generally have heterozygous mutCALR clones and some residual wild-type hematopoiesis. To explore whether RPG4 CAR T cells could identify and kill mutCALR cells when present at both low and high frequencies, RPG4 CAR T cells were tested against mixtures of mutCALR+ and wild-type cells. RPG4 CAR T cells depleted 96.7% of mutCALR+ cells when present at only 10% frequency among wild-type hematopoietic cells (Fig. 3C). Quantifying the mutant clone burden (variant allele frequency, VAF) before and after CAR T cell co-culture showed a substantial reduction in VAF ranging between -20 to -79%, only in mutCALR+ samples (Fig. 3D), reflecting 50-75% clearance of the mutant clone stem cells, correlating with the baseline mutCALR burden (Fig. 3E).

### Eltrombopag boosts CAR T killing of low target-expressing cells

Progression to AP/BP-MPN is associated with a particularly poor outlook (survival ∼ 6 months) and no effective therapeutic options (*21*). We noted that RPG4 CAR T cells demonstrated poor cytotoxicity when tested against CD34+ cell samples from patients with AP/BP-MPN (Fig. 3F). As leukemic transformation of MPN is typically myelogenous rather than megakaryoblastic in phenotype (*22*), we speculated that the reduced killing may be due to lower cell surface expression of TpoR, and therefore mutCALR, on leukemic blasts. In keeping with this, we detected lower TpoR expression on AP/BP-MPN samples that were ‘poor responders’ to the CAR T therapy *in vitro* than samples from patients who were ‘responders’ (Fig. 3G). We hypothesized that pharmacological TpoR stimulation may promote mutCALR expression by promoting TpoR expression and/or driving megakaryocytic differentiation of blasts. Several TpoR agonists are currently in clinical use for the treatment of thrombocytopenia, including small molecule agonists (e.g. eltrombopag) and peptibody therapies (e.g. romiplostim). We evaluated the impact of recombinant human Tpo (rhTpo), eltrombopag and romiplostim on cell surface TpoR expression of Marimo-TpoR cells. While rhTpo drove internalization of the TpoR, eltrombopag markedly increased TpoR expression (Fig. 3H), and this corresponded with increased mutCALR CAR T cell mediated killing of both cell lines and samples from patients with AP/BP-MPN (Fig. 3I and fig. S6a-c).

### Comparison of CAR T cell cytotoxicity to JAKi therapies

JAK inhibitor therapies are frequently used in myelofibrosis to control symptoms, blood counts and splenomegaly. These agents are non-selective for the mutant clone and generally have minimal impact on the VAF in myelofibrosis (23). To benchmark the cytotoxic efficacy of the mutCALR CAR T cell therapy to current therapies, we compared the CAR T therapy to ruxolitinib and momelotinib – agents that are used as first-line treatments for myelofibrosis. CAR T cell therapy showed significantly greater cytotoxicity than either JAKi (p<0.001, Suppl. fig. S6d).

### Evaluation of CAR T cell function in a 3D human multi-cellular tissue environment

Simple co-culture assays do not determine how CAR T cell function is influenced by the tumor microenvironment. This is particularly important to evaluate in myelofibrosis due to the prominent fibrosis, inflammation and abundance of immunoregulatory cytokines (*24*). We recently developed a human bone marrow organoid model in which multi-cellular hematopoietic niches are generated from human induced pluripotent stem cells (iPSCs) (*25*)(*26*). The organoids undergo cancer cell-induced niche remodeling after seeding with HSPCs from myelofibrosis patients, and addition of TGFβ can also induce organoid fibrosis, enabling evaluation of novel therapies in a relevant human tissue context.

As human bone marrow organoids have not previously been used to evaluate CAR T cell therapies, we first confirmed that CAR T cells could engraft and remain viable in the organoids. Immunofluorescent imaging and flow cytometry demonstrated that CAR T cells rapidly engrafted into the organoids, being clearly visible throughout the organoid body and vessels by 6-12 hours and persisted for up to 7 days (Fig. 4A). Cell type abundance and viability of organoid stroma was unchanged following addition of CAR T cells, excluding a significant allo-reactive effect (fig. S7a & S7b). To initially test if CAR T cells were functional within organoids, CD19 or RPG4 CAR T cells were added to organoids engrafted with either SEM cells (a B-cell leukemia cell line), or Marimo-TpoR cells. Both CAR T cells selectively depleted cognate cell lines (Fig. 4B).

**Fig. 4.**
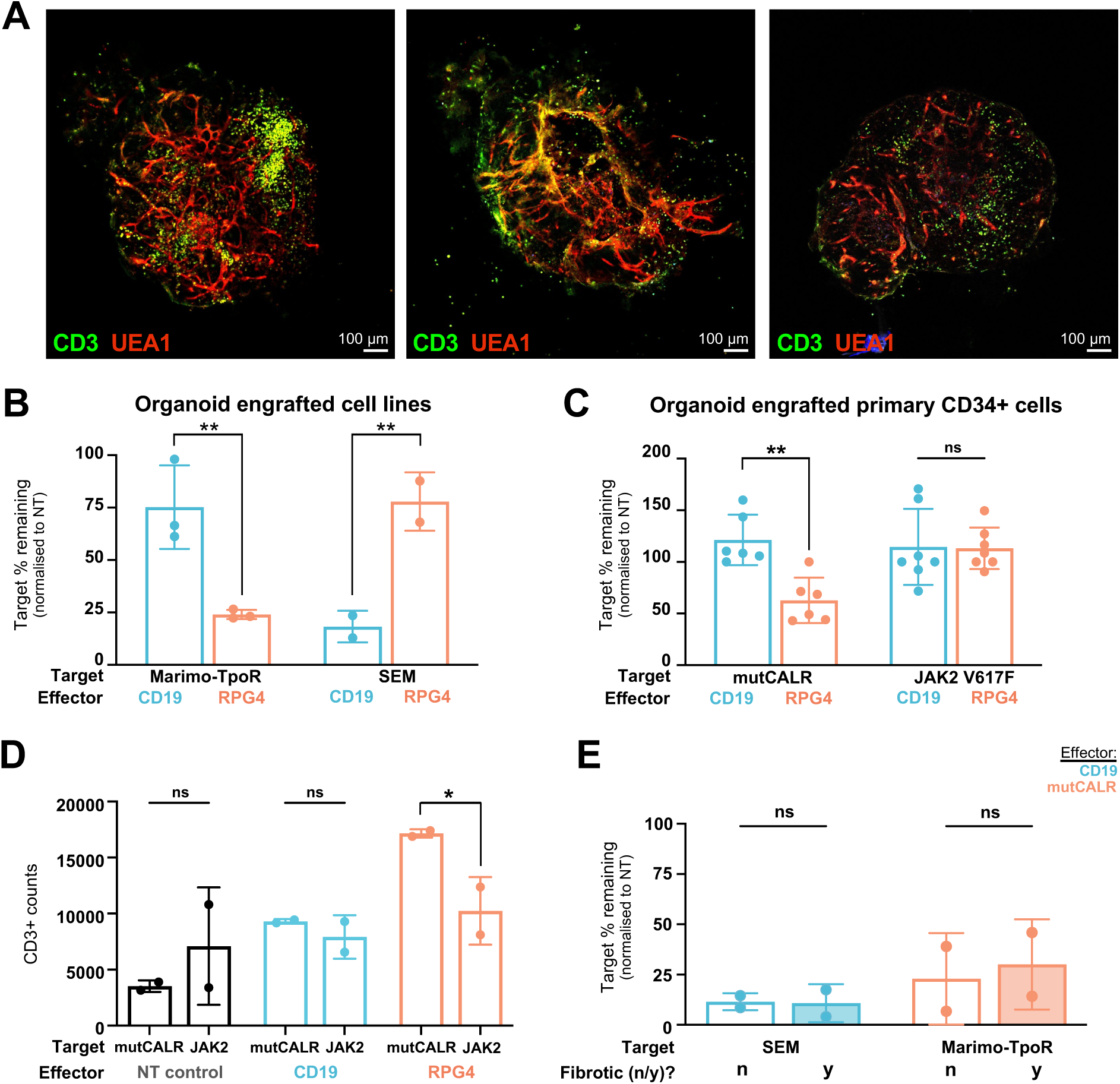
mutCALR-directed CAR T cells specifically target malignant cells in a 3d marrow organoid model. **(A)** Representative images of organoids engrafted with CAR T cells confirming engraftment and dissemination of CAR T cells throughout the organoid parenchyma, associating with the organoid vessels. Green – CD3 (CAR T cells); red – UEA1 (vessels). **(B)** % target cells remaining (normalised to non-transduced [NT] control condition) in organoids following engraftment with either mutCALR+ Marimo cells expressing the thrombopoietin receptor (TpoR) or CD19+ SEM cells, followed by addition of either RPG4 CAR T cells or CD19-targeting CAR T cells. Each dot represents n = 4 organoids, for 2-3 independent experiments. **(C)** % target cells remaining (normalised to NT control) in organoids engrafted with CD34+ hematopoietic stem/progenitor cells from patients with either mutCALR+ (n = 6) or JAK2V617F+ (n = 7) myelofibrosis, followed by addition of mutCALR CAR T cells or CD19 targeting CAR T cells as a control. **(D)** Number of CD3+ events (representing CAR T cells) in organoids engrafted with either mutCALR+ or JAK2V617F+ cells plus either NT, CD19- or mutCALR-targeting CAR T cells. Each dot represents n = 4 organoids in 2 independent experiments. **(E)** % target cells remaining after addition of either CD19- or mutCALR-targeting CAR T cells and cognate CD19+ or mutCALR+ target cell line in ‘healthy’ or ‘fibrotic’ (TGFβ pre-treated) organoids. TGFβ is washed out after the induction of fibrosis prior to engraftment with target cells and CAR T cells. Comparisons made using 2-way ANOVA. * (p<0.05), ** (p<0.01), *** (p<0.001), ****(p<0.0001).

CAR T cells were next tested against organoids engrafted with HSPCs from patients with myelofibrosis. CAR T cells were added 24 hours after the myelofibrosis HSPCs, to allow the cancer cell cells to home to and engraft the organoids. Cytotoxicity was evaluated 72 hours after the addition of CAR T cells. CAR T cells exhibited selective killing of mutCALR+ but not JAK2V617F+ HSPCs (Fig. 4C) with CAR T cell expansion only in response to mutCALR targets (Fig. 4D).

Next, we tested CAR T function against mutCALR+ target cells engrafted into organoids that were pre-treated with TGFβ to induce fibrosis (collagen 1 deposition and alpha smooth muscle actin expression, fig. S7c), followed by wash-out of TGFβ, to examine killing efficacy in a fibrotic tumor environment. CAR T cells were able to engraft the fibrotic niches (fig. S7d) and demonstrated excellent cytotoxicity, confirming effective penetration and efficacy in fibrotic marrow tissue (Fig. 4E).

### Functional and molecular consequences of CAR T mediated cancer cell killing

The human organoid model provided a unique opportunity to examine the functional impact and molecular consequences of RPG4 CAR T cells on healthy and malignant hematopoietic cells when co-engrafted within the MPN bone marrow microenvironment. Single cell RNA sequencing (scRNAseq) was performed on human organoids engrafted with CD34+ HSPCs from a patient with high-risk mutCALR+ myelofibrosis (table S1), with and without addition of mutCALR CAR T cells generated from a healthy donor and engrafted 24 hours later. In this system, the hematopoietic niche, myelofibrosis cells and CAR T cells each derive from different donors, therefore we were able to genetically de-multiplex each cellular compartment using single nucleotide polymorphisms unique to each donor (Fig. 5A, fig. S8a) (*27*). After QC, 35,794 cells were included in the analysis, including 15,164 iPSC-derived organoid stromal cells, 13,810 myelofibrosis cells and 6,820 CAR T cells. Cell types were annotated using canonical marker genes (Fig. 5B).

**Fig. 5.**
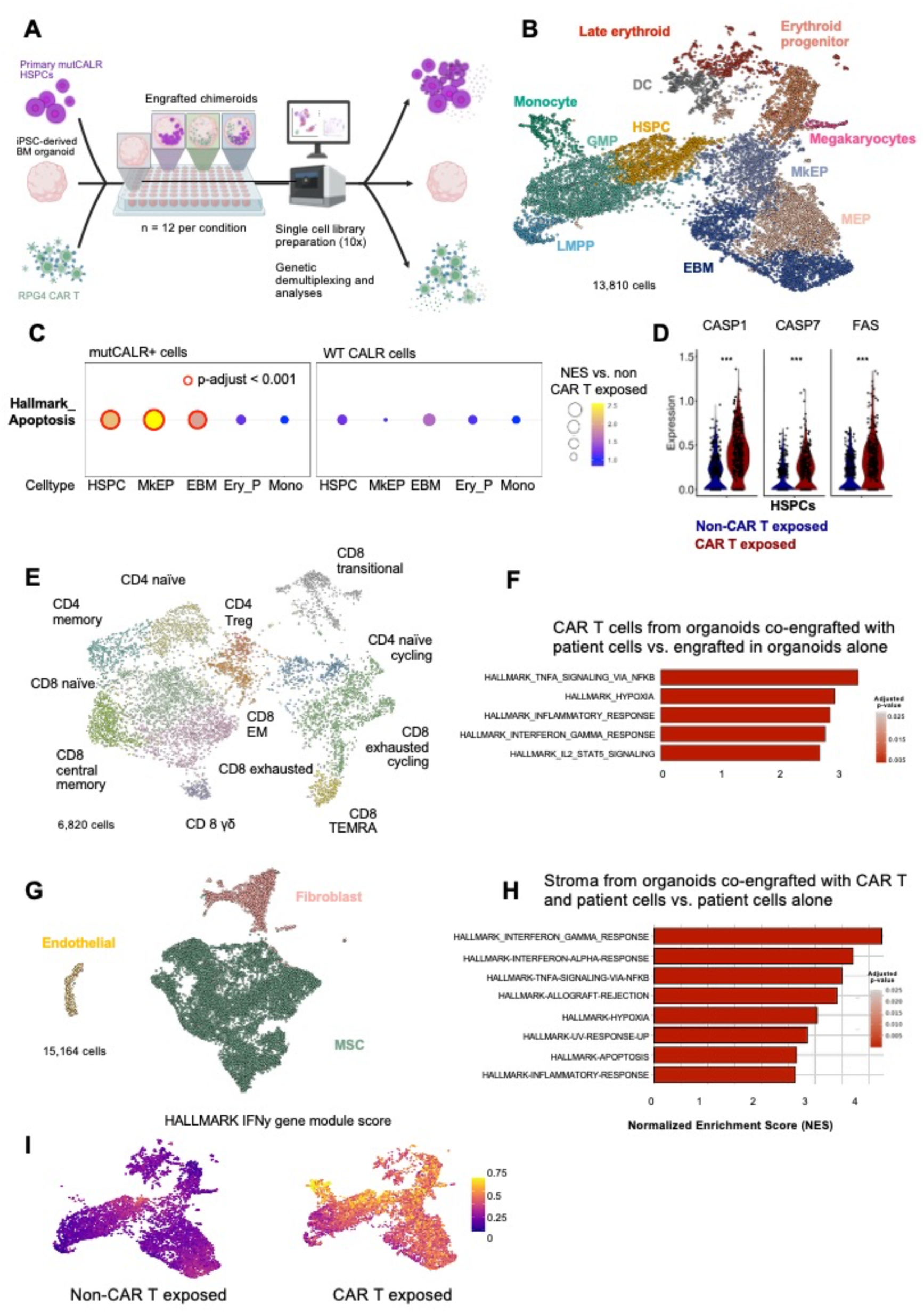
Cellular and molecular impact of mutCALR CAR T cell targeting in 3D human tissue environment. **(A)** Schematic illustrating ‘chimeroid’ model in which organoids are engrafted with CD34+ stem/progenitor cells from a patient with mutCALR+ myelofibrosis, healthy donor-derived mutCALR CAR T cells, both or neither. Organoids were analysed by single cell RNAseq and the use of different donors enables genetic de-multiplexing of iPSC-derived organoid cells, myelofibrosis cells and CAR T cells for detailed interrogation. (B) Uniform manifold approximation and projection (UMAP) plot of cells derived from the donor with myelofibrosis (13,810 cells), showing cluster annotation. (C) Expression of the HALLMARK_APOPTOSIS geneset in mutCALR+ (left) and wildtype (WT, right) cells exposed to RPG4 CART compared to cells not exposed to CAR T cells. Normalised enrichment score (NES) is shown, with significant enrichment of apoptosis only in mutCALR+ cells that express TPO-R (HSPCs, MkEP, EBM). Genotype (Type 2 mutCALR vs WT) was assigned using VarTrix. (D) Expression of CASP1 (log2FC 1.46), CASP7 (log2FC 1.05) and FAS (log2FC 1.3) in mutCALR+ HSPCs exposed to RPG4+ CAR T cells (red) vs non-CAR T cell exposed (blue). *** adjusted p-value < 0.001. (E) UMAP showing induction of inflammation across all wild type and mutant MPN cell types after CART exposure (HALLMARK IFNy gene module score). (E) UMAP of CAR T cells isolated from control organoids or organoids engrafted with myelofibrosis cells showing annotation of T cell subtypes. (F) Gene set enrichment analysis comparing gene expression profiles of CAR T cells extracted from myelofibrosis-engrafted organoids (n = 12) vs. organoids without myelofibrosis cells. (G) UMAP of iPSC-derived stromal cells, showing cluster annotations. (H) Gene sets enriched in organoid stroma cells after co-engraftment with CAR T cells and primary mutCALR+ cells. (I) IFNγ gene module score in myelofibrosis cells from organoids co-engrafted with CAR T cells vs. non-CAR T cell exposed. Abbreviations: MEP – megakaryocyte-erythroid progenitor cells; MkEP – Megakaryocyte and Erythroid progenitors; EBM – eosinophil-basophil-mast cell progenitors; HSPC – hematopoietic stem/progenitor cells; LMPP – lympho-myeloid primed progenitor cells; GMP – granulocyte-monocyte progenitor cells; MSC – mesenchymal stromal cells).

We first extracted the myelofibrosis cells for detailed analysis. Previous studies have shown that MPN driver mutations induce myeloid-biased hematopoiesis, with an expansion of megakaryocytes (*28*)(*29*), as well as basophil and mast cell progenitors (*24*). In keeping with this, analysis of the patient-derived cells captured from the organoids showed prominent clusters of megakaryocyte and basophil/mast cell progenitors, confirming that the intrinsic skewing of hematopoiesis in myelofibrosis was sustained within organoid niches (Fig. 5B). Variant calling was performed on the patient cells to distinguish mutant CALR and wild-type patient cells. Genotypes were assigned to 79.4% (9862/13810) cells (fig. S9a), while it was not possible to assign genotypes in 28.6% (3948/13810) cells, likely due to low expression of CALR or low coverage of that region (fig. S9b). Analyzing the differential gene expression in mutCALR+ and wild-type cells from organoids co-engrafted with CAR T cells compared to cells from organoids with no CAR T cells revealed significant enrichment of apoptosis pathway genes selectively in cellular subsets known to express TpoR, including HSPCs, megakaryocyte/erythroid progenitors (MkEP) and eosinophil-basophil-mast cell progenitors (EBM) (Fig. 5C). No enrichment of apoptosis pathway genes was observed in TpoR negative mutant cells or TpoR+ wildtype cells, confirming on-target selectivity in primary cells in a relevant human tissue model. Consistent with the pathway-level enrichment, gene-level analysis showed significant upregulation of key apoptotic mediators including CASP1, CASP7, and FAS within these targeted mutCALR⁺ populations following CAR T cell exposure (Fig. 5D, p < 0.001). Higher MPL expression was detected in myelofibrosis cell types that were targeted by the mutCALR CAR T cells (HSPCs, MkEP and EBM) than cell types that were not targeted following addition of CAR T cells (fig. S10a). These findings indicate that RPG4 CAR T cell activity selectively induces apoptosis in mutant cells that express the mutCALR-TpoR complex.

Examining the organoid-engrafted CAR T cells, subsets of CD8 naïve, CD8 effector memory, CD4 naive and CD4 memory as well as regulatory and γδ T cells were identified (Fig. 5E). The differential abundance and transcriptional phenotypes of CAR T cells engrafted into organoids alone vs. engrafted into organoids containing mutCALR+ myelofibrosis cells were examined. This revealed strong enrichment of activation markers and inflammatory signaling pathways including TNFα, IFNγ and IL2-STAT5 in CAR T cells that had been exposed to mutCALR target cells (Fig. 5F), and a marked expansion of CD4 and CD8 naïve T cell subsets (fig. S10b).

Finally, we examined the impact of CAR T cell mediated target killing on the marrow stroma and bystander hematopoietic cells. Prolonged cytopenias are frequently observed after CAR T cell therapy, which has been attributed to inflammatory damage to the hematopoietic niche (*30*), although this is challenging to study mechanistically in patients. For example, a study of 21 patients receiving CAR T cells for B cell malignancies reported a significant decrease in CD271+ mesenchymal stromal cells (MSCs) identified on bone marrow biopsies, particularly in those with prolonged cytopenia (*31*). In keeping with this, we found a significant enrichment of IFNγ, IFNα, TNF signalling and apoptosis pathway gene sets in stromal cell subtypes from organoids co-engrafted with CAR T cells and patient cells compared to organoids engrafted with patient cells alone (or CAR T cells alone), reflecting an inflammatory response to CAR T cell cytotoxicity (Fig. 5G, H). Mirroring the inflammation in the stroma, there was a global increase in inflammatory response genes associated with IFNγ signalling in all cell types from organoids engrafted with both CAR T cells and primary cells regardless of mutant status, indicating an inflammatory response in bystander cells including non-malignant populations in response to CAR T cell activation (Fig. 5I).

### mutCALR CAR T cell therapy can control the growth of mutCALR+ leukemia in vivo

To evaluate the in vivo efficacy of RPG4 CAR T cells, we used two human xenograft models, engrafting human acute myeloid leukemia cell lines Molm-14 and Marimo into immunodeficient mice. Both cell lines induce a leukemia-like disease with widespread, aggressive cell growth with high signal in the bone marrow. Molm-14 cells were originally derived from a patient with an aggressive myeloid leukemia driven by FLT3-ITD and MLL-AF9 gene fusions. As these cells do not harbor a CALR mutation nor express cell surface TPO-R, they were transduced with a construct encoding the del52 mutCALR C-terminus fused to a CD8 stalk (mutCALR-CD8STK) to achieve mutCALR cell surface expression, and with firefly luciferase (Fluc) to enable monitoring of tumor burden via bioluminescence Imaging (BLI, Fig 6A). RPG4 CAR T cells resulted in effective control of leukemic growth, as assessed by BLI (Fig 6B), and led to a significant prolongation of survival compared with mice treated with the negative control anti-CD19 CAR T cells (log-rank p = 0.0012, Fig 6C).

**Fig. 6.**
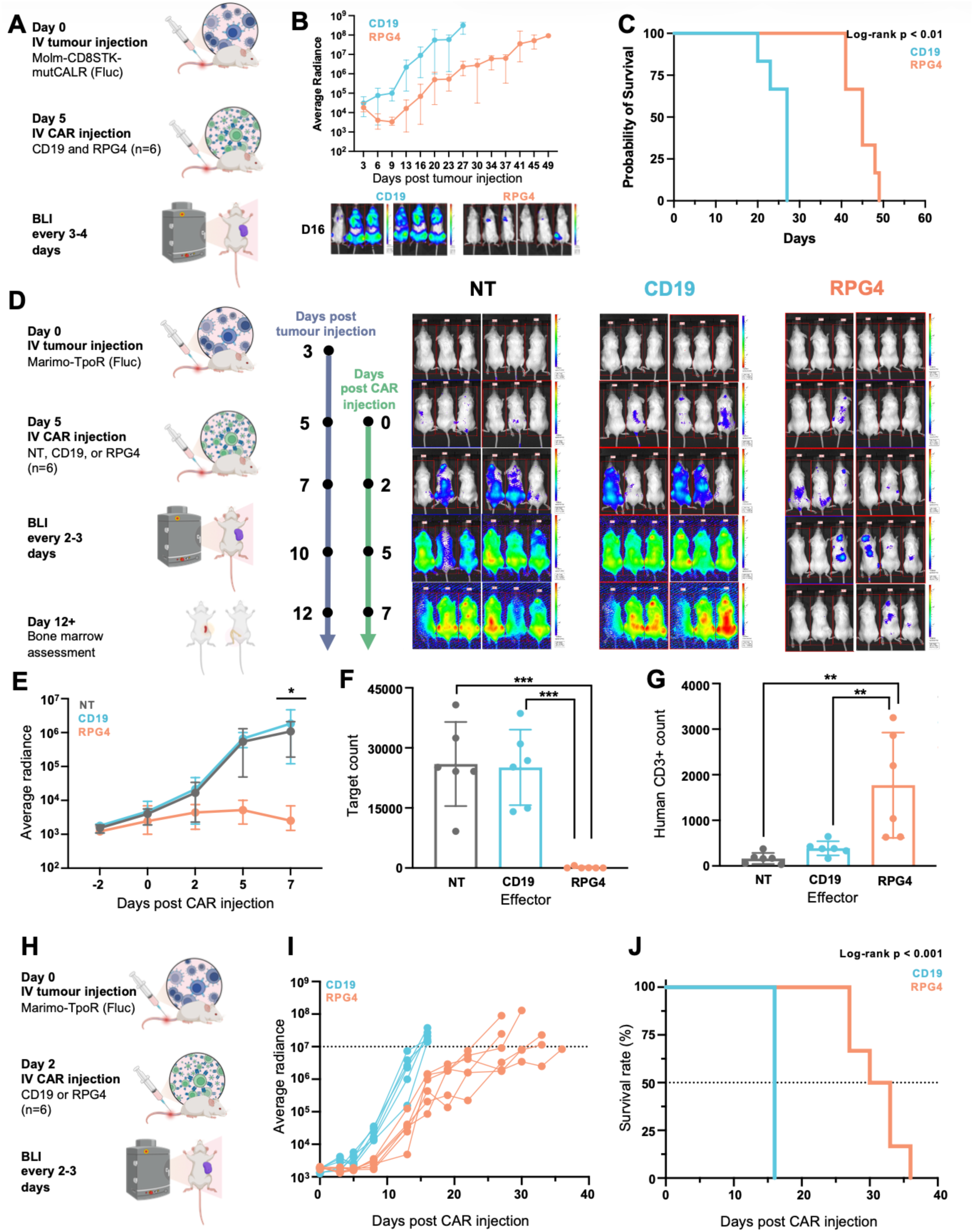
RPG4 CAR T cells control disease and improve survival in a mutCALR *in vivo* leukemia xenograft model. **(A)** Schematic overview of the xenograft leukemia model using the Molm-14 cell line transduced with CD8-Stalk with mutCALR C terminus and fluorescent reporter. **(B)** Quantification of average radiance indicating disease burden for each treatment group. **(C)** Kaplan–Meier survival analysis comparing mutCALR CAR T–treated mice with those receiving CD19 CAR T cells (as a negative control). **(D)** Left - schematic overview of the xenograft leukemia model. Right – bioluminescence imaging (BLI) of mice at indicated time points. **(E)** Quantification of average radiance in each treatment group over time, indicative of leukemic burden. **(F)** Number of BFP+ leukemia cells in the bone marrow/1E6 live cells, quantified by flow cytometry for BFP+ cells. **(G)** Number of human CD3+ T cells/1E6 live cells in the bone marrow quantified by flow cytometry. **(H)** Schematic overview of the survival cohort. **(I)** Quantification of average radiance in individual mouse over time. **(J)** Kaplan–Meier survival analysis comparing mutCALR CAR T–treated mice with those receiving CD19 CAR T cells (as a negative control). 2-way ANOVA was used for comparisons between groups in bar charts, and Kaplan-Meier analysis for the survival differences. ** (p<0.01), *** (p<0.001).

RPG4 CAR T cells were also tested in xenografts using the Marimo-TpoR cell line, which derives from a patient with acute myeloid leukemia transformed from mutCALR+ Essential Thrombocythemia. Two days after tumor cell inoculation, mice were injected with 5 x 10^6^ mutCALR-targeting CAR T cells or controls (non-transduced human T-cells or anti-CD19 CAR T cells, Fig. 6D). Administration of mutCALR CAR T cells led to a significant reduction in leukemic burden compared to CD19-targeting and non-transduced T cells (P < 0.05, Fig. 6D & 6E). Leukemic cells were almost completely eliminated from the bone marrow of mice, with significant human T-cell expansion (Fig. 6F & 6G).

A survival analysis was then performed in a separate cohort of mice (Fig. 6H). The disease progression was monitored via bioluminescent (BLI) imaging and pre-set sacrifice criteria were established upon reaching an average radiance of 1E7 photons/s/cm²/sr. Treatment with mutCALR CAR T cells led to control of the tumor growth and significant prolongation of survival (Fig. 6I & 6J, p=0.009). Control mice had rapidly progressive disease with significant luminescence signal in the bone marrow, while the mutCALR group achieved a >2-fold longer survival (36 days vs 16 days, Fig. 6I).

## DISCUSSION

Currently approved therapies for MPNs include cytoreductive agents and JAK inhibitors – treatments that may control myeloproliferation and alleviate symptoms, but are not curative, and almost all patients eventually become refractory to therapy. Agents targeting pathways beyond JAK are being explored, however, none of the licensed therapies exert selective control of mutant HSCs. To-date, allogeneic HSC transplant remains the only intervention that can substantially improve long-term outcome, although only a small minority of patients benefit (*32*) and it is associated with significant toxicities including up to 40% transplant-related mortality. A therapeutic that potently and selectively depletes HSCs bearing the driver mutation is required, to allow healthy hematopoiesis to recover and predominate.

The unusual biology behind mut*CALR*-driven MPNs leaves a mutated CALR protein with a common nonsense carboxy terminus attached to the TpoR on the surface of malignant cells. This provides an opportunity for selective, immune-mediated depletion of disease-driving HSCs.

CAR T cells can achieve lasting remissions, including in patients with cancers resistant to standard chemotherapy and therapeutic antibodies (*33*). CAR T cells may be more potent than other approaches – for example, a recent meta-analysis showed that CAR T therapy led to more frequent durable remissions than TCEs in patients with diffuse large B cell lymphoma (*15*). Additionally, CAR T cell therapies can be further augmented by additional engineering (*34*) e.g. by expression of a dominant-negative TGFβ receptor to improve function in a challenging, TGFβ-rich tissue context such as myelofibrotic marrow (*35*), or with other genetically encodable modifications (*34*).

In this paper we describe RPG4, a scFv which binds selectively to the nonsense frameshift sequence at the C-terminus of mutCALR. RPG4 as an scFv-Fc format only bound cells expressing mutCALR complexed with TpoR on the cell surface. CAR T cell recognition of TpoR/mutCALR with a standard CAR is not a given since the target antigen is compound, and spans two sizeable proteins which may prevent immune synapse formation and full CAR T cell activation (*36*). Despite the complex nature of the target antigen, RPG4 CAR T cells demonstrated highly selective and potent killing against cell lines and primary cells that co-express both the TpoR and mutCALR. Crucially, when tested on HSPCs from patients with MPN, RPG4 CAR T cells induced a marked reduction in mutant clone burden in mutCALR samples, while showing no cytotoxicity against JAK2 V617F samples. Although promising efficacy of RPG4 CAR T cells was observed in human organoids and in cell line xenograft studies, these experiments may not reflect clinical activity and future studies are required to determine whether the RPG4 CAR T cell therapy can meaningfully control disease with an acceptable toxicity profile in patients.

Patients whose disease progresses to AP/BP-MPN have a particularly poor outlook (survival ∼ 6 months) and no effective therapeutic options (*21*). We noted poor cytotoxicity against samples from patients with AP/BP-MPN. As leukemic transformation of MPN is typically myelogenous rather than megakaryoblastic in phenotype (*22*), we speculated that the reduced killing was due to lower cell surface expression of TpoR. We hence explored increasing TpoR expression through pharmacological TpoR stimulation. Unexpectedly, we found that while rhTPO drove internalization of the TpoR, the small molecule TpoR agonist eltrombopag markedly increased TpoR expression with corresponding increased killing by RPG4 CAR T cells. Finally, we demonstrated potent clearance of mutCALR malignancies in a 3D, multicellular tissue environment – both *in vitro* in a bespoke human bone marrow organoid model, and *in vivo*.

Several key questions require further study. Firstly, the depth of molecular response that is required for meaningful disease remissions in patients with MPNs is not known. *CALR* mutations confer only a modest competitive advantage for the malignant HSCs, and clonal expansion occurs over decades (*37*). Therefore, a reduction in the mutant clone burden without complete (or deep) molecular remission may be clinically meaningful and sufficient to impede progression. Secondly, the ability of the non-malignant HSCs to restore heathy hematopoiesis after CAR T therapy in myeloid malignancies remains unclear.

Collectively, the data presented here highlight a new therapeutic strategy for patients with MPNs. Exploiting the unique biology of mutCALR, CAR T cells can be directed to selectively deplete the cancer stem cells. This sets the stage for use of CAR T cell therapy as molecularly targeted cancer therapy – specifically targeting a disease-driving cancer neoantigen that is the direct product of an oncogene. While future clinical studies are required, this approach has potential to improve clinical outcomes for patients with mutCALR-driven cancers. In addition, we advanced a human bone marrow organoid model that permits detailed characterization of CAR T cell function and the cellular/molecular consequences on the cancer clone in the relevant human TME, including inflammatory responses in hematopoietic niche cells. This offers a powerful approach for the evaluation of immunotherapies targeting blood cancers beyond MPNs, that may accelerate successful clinical translation.

## Funding

Medical Research Council UK MR/W000326/1 (AR)

Blood Cancer UK 23005

National Institutes of Health US graduate partnership program (ZCW)

National Institute of Health Research UK

Oxford Biomedical Research Centre UK

Cancer Research UK Senior Fellowship (BP)

Rosetrees Trust (BP)

Ludwig Cancer Research Institute (BP)

## Author contributions

AR, MP, CR and JL conceived the project and acquired funding. AR, ZCW and IG performed the *in vitro* experiments. ZCW performed the computational analysis for the scRNAseq. AB and MF performed the surface plasmon resonance for the binders. IG and AR performed the *in vitro* experiments for binder characterisation and CAR T validation against cell lines, while ZCW performed the *in vitro* experiments against primary samples AR performed the *in vivo* experiments with the contribution of IG. YS, ARR, AAO provided input for the organoid experiments. ChB performed and supervised computational analysis and contributed to writing the manuscript. CaB provided input for the *in vivo* experiments. GW and MM provided input for the CAR T experiments. DR, SK and MRJ performed the histopathology staining. IH helped with the conception of the TpoR boosting idea. AC collected the patient samples used and NH assisted with sample processing. AK provided overall supervision and input for the organoid experiments. LM and AK provided input and assistance with immunofluorescence imaging. AM provided input throughout the project. MP, CR and BP provided regular input and supervision on the conduct of experiments. BP designed and supervised the experiments involving primary patient cells and organoid CAR T testing and co-wrote the manuscript. The remaining experiments were designed and conceptualized by AR, IG, ZCW, BP and MP. All authors read and approved the submitted manuscript.

## Data and materials availability

All data associated with this study are present in the paper or the Supplementary Materials. All plasmids used in the study including CAR T constructs and mutCALR expression vectors are available from the first author, AR, upon request and completion of a material transfer agreement. All raw and processed sequencing data generated in this study have been submitted to the NCBI Gene Expression Omnibus.

---

**S1.**
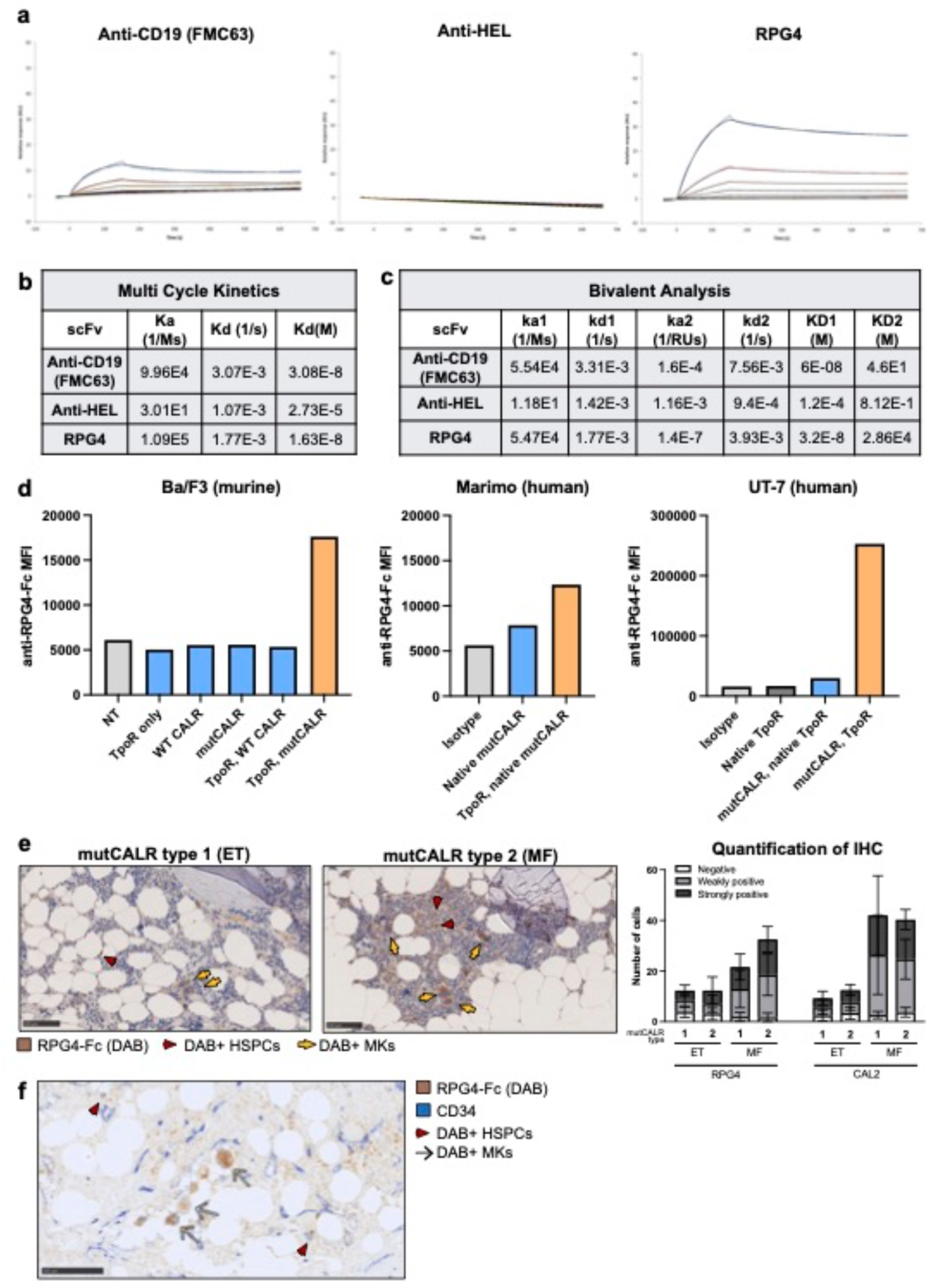
RPG4 binding kinetics to purified mutCALR C-terminus and bone marrow biopsies. **(A)** Surface plasmon resonance binding kinetics showing response (relative units, RU) of RPG4 (right) vs. FMC63 (anti-CD19; left) and anti-HEL (Hen egg lysozyme; middle) binders as relevant and internal controls respectively. Analytes were injected at variable concentrations with contact time 150s and dissociation time 500s at a flow rate of 30 µl/min. Response is shown in relative units (RU); increasing concentrations are represented by ascending curves. **(b, c)** Binding kinetics for each scFv in response to mutCALR C-terminus protein. **(d)** Quantification of mean fluorescence intensity (MFI) of anti-RPG4-Fc binding to cell lines **(e**) Representative immunohistochemistry staining (left) and formal analysis (right) using RPG4 scFv on bone marrow biopsies from 4 patients with either essential thrombocythemia (ET) or myelofibrosis (MF) driven by Type 1 (N=2) or Type 2 (N=2) *CALR* mutations. Brown colour (3, 3’-diaminobenzidine, DAB) indicates positive staining. Arrows indicate DAB+ megakaryocytes (MK) (yellow) and HSPCs. **(f)** Representative bone marrow biopsy section co-stained for CD34 (blue) and RPG4-Fc (brown). Red arrows indicate CD34+ mutCALR+ hematopoietic stem/progenitor cells (HSPCs); white/black arrows indicate CD34- mutCALR+ megakaryocytes (MK).

**S2.**
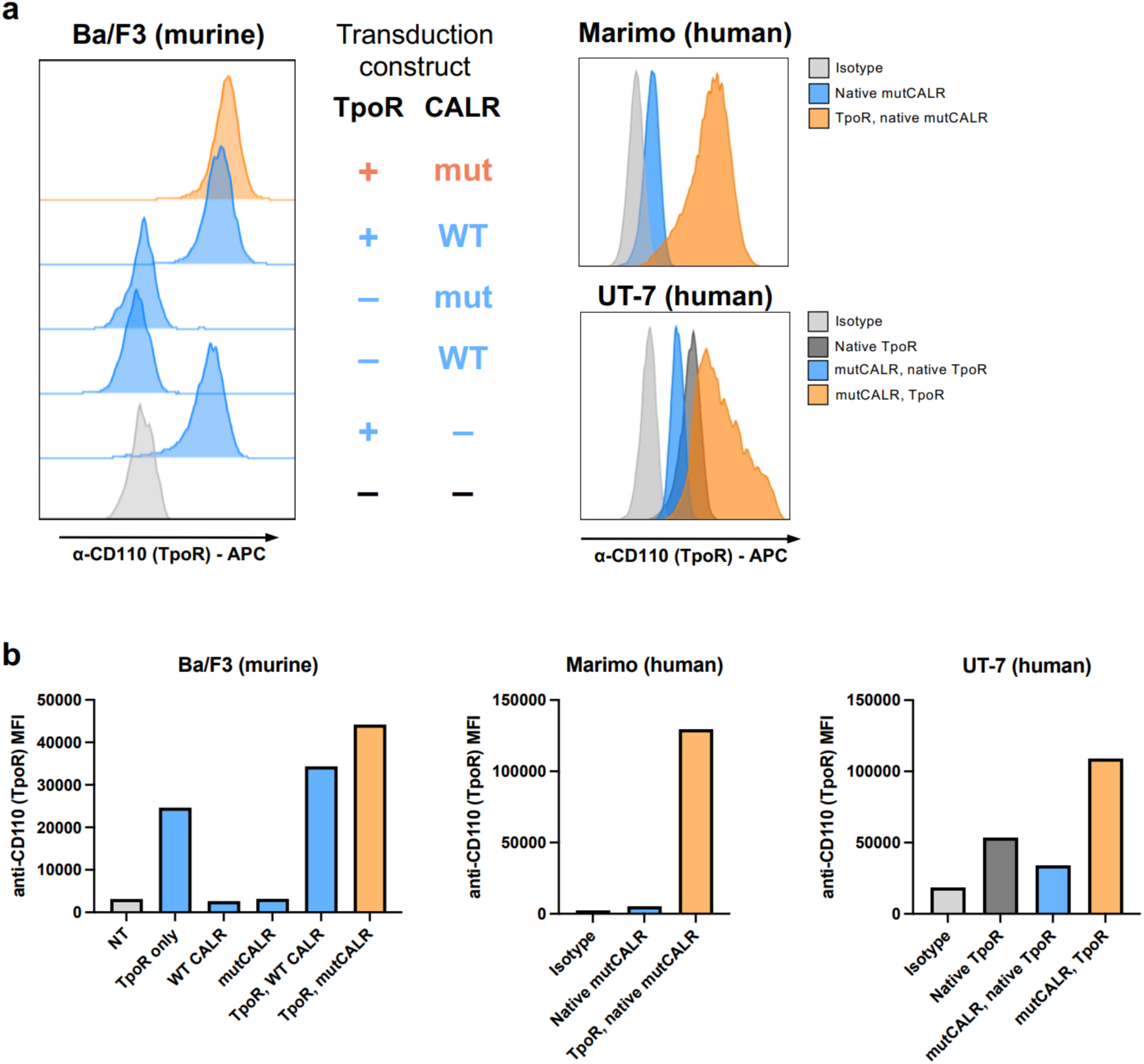
Expression of the thrombopoietin receptor (TpoR) on panel of engineered cell lines. **(A)** Mean fluorescence intensity (MFI, normalised to mode) of TpoR expression of murine pro-B (Ba/F3) cell line (left) including non-transduced cells (NT) and cells engineered to express TpoR alone, and also with wildtype (WT) or mutant (mut) CALR as indicated. TpoR expression of engineered human cell lines including mutCALR+ acute myeloid leukemia Marimo cell line (top right) transduced with/without TpoR, and megakaryoblastic UT-7 cells (bottom right) transduced with mutCALR with/without TpoR. Isotype control antibody also shown in light grey. **(b)** Charts showing MFI of TpoR expression for each cell line. Representative analyses shown.

**S3.**
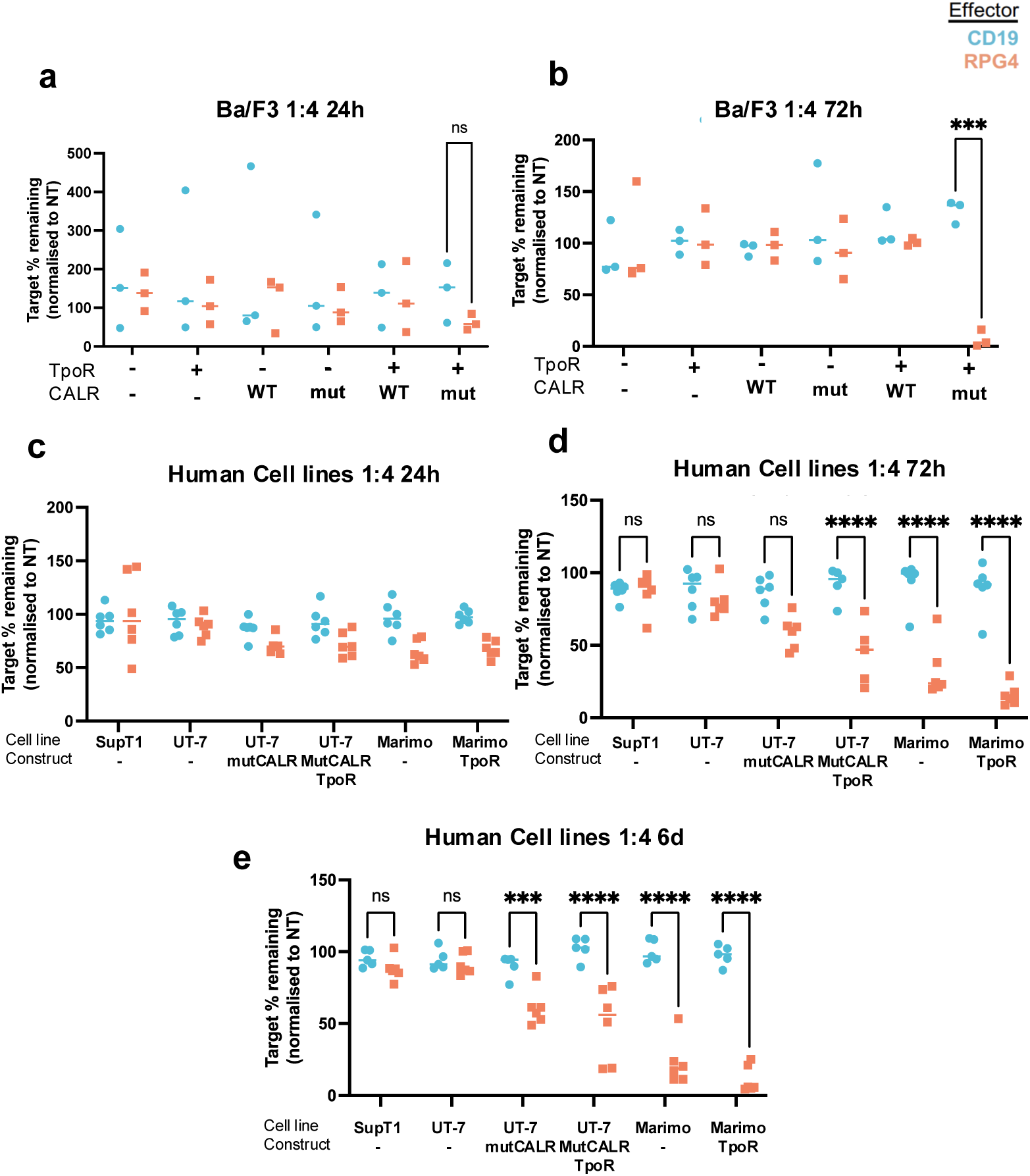
MutCALR targeting RPG4 CAR T cells eliminate target bearing cell lines at low effector:target (ET) ratios. **(a, b)** Flow cytometry-based killing assays using RPG4 CAR T cells against target positive and negative Ba/F3 cells in an E:T ratio of 1:4 and readouts at **(a)** 24 hours (hrs) and **(b)** 72 hrs. Cell numbers were normalised to the condition where non-transduced (NT) T cells were added. CAR T cells were generated from n=3 healthy donors, and human CD19-targeting CAR T cells were used as a negative control. **(c – e)** Killing assay of target negative cells (SupT1, UT-7-NT), low-expression (UT-7 mutCALR, Marimo NT) and high-expression (UT-7 mutCALR/TpoR, Marimo-TpoR human cell lines using RPG4 CAR T cells generated from healthy donors (n=6 donors). CD19 targeting CAR T cells were used as a negative control. E:T ratio was 1:4 and readouts shown for **(c)** 24 hrs, **(d)** 72 hrs and **(e)** 6 days.

**S4.**
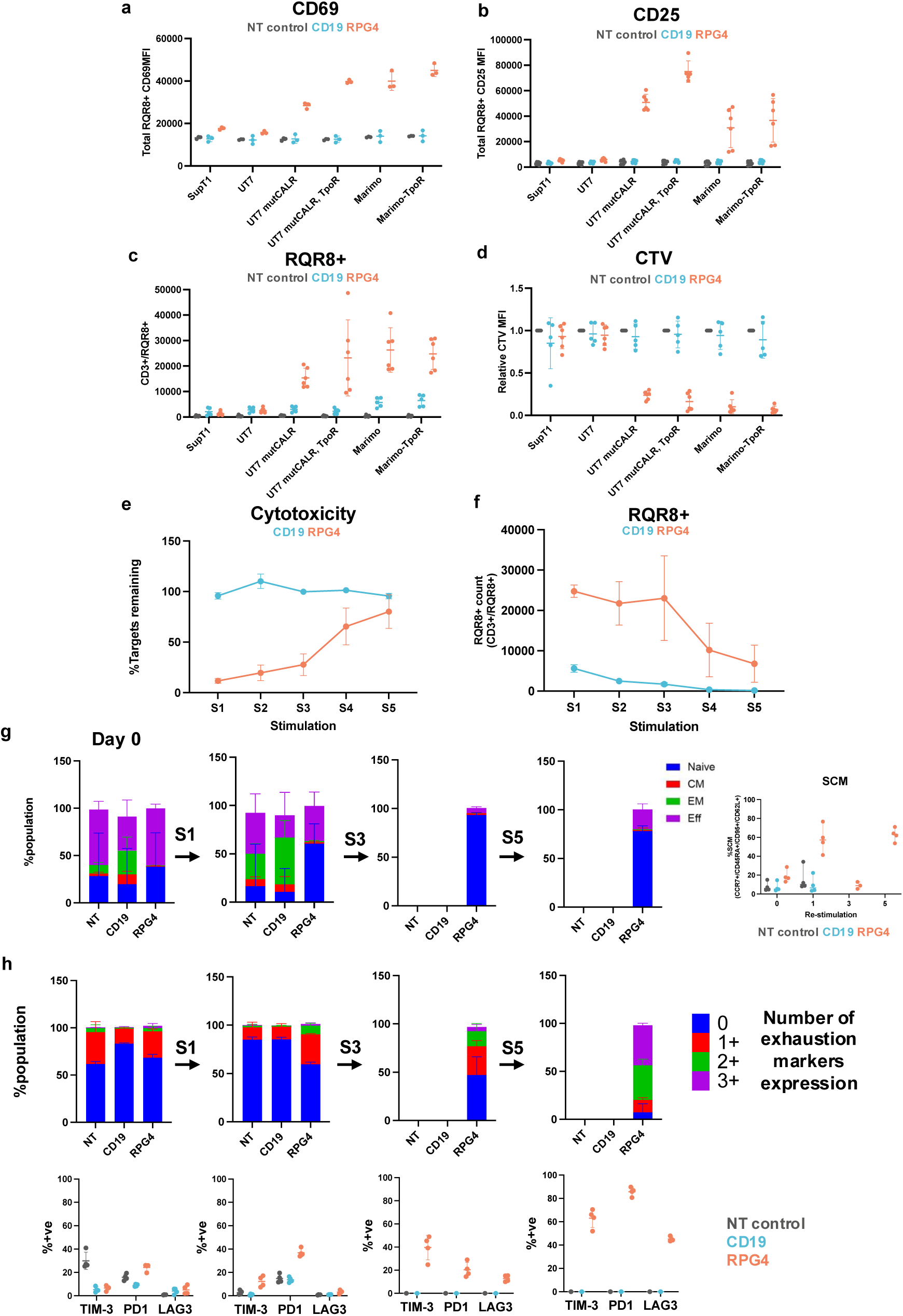
Activation and co-stimulation assays for mutCALR-targeting RPG4 CAR T cells. **(a, b)** Mean fluorescence intensity (MFI) of expression for **(a)** CD69 and **(b)** CD25 for RPG4 CAR T cells at 24h and 72h of co-culture with target negative (SupT1, UT-7-NT), low-expression (UT-7 mutCALR, Marimo NT) and high-expression target cells (UT-7 mutCALR/TpoR, Marimo- TPOR) (N=3). **(c, d)** CAR T cell proliferation assessed by **(c)** quantification of RQR8+CD3+ events and **(d)** Cell trace violet (CTV) stain dilution following 6 days of co-culture with the target cells as indicated**. (N=3) (e)** Percentage of target cells remaining and **(f)** RQR8+CD3+ CAR T cell counts relative to non-transduced (NT) T cell condition after 5 rounds of stimulation with Marimo**-T**poR cells (S1-5) (N=3). **(g)** T-cell phenotypes and **(h)** expression of exhaustion markers TIM3, PD1, LAG3 following each round of stimulation (N=3). T-cell phenotypes were assigned as follows Naive T-cells: CCR7+ CD45RA+, Central Memory T-cells (CM): CCR7+, CD45RA-, Effector Memory T-cells (EM): CCR7-, CD45RA-, Effector T-cells (EFF): CCR7-, CD45RA+, and stem like memory (SCM) CCR7+ CD45RA+ CD95+ CD62L+. Exhaustion markers included Lag3, TIM3 and PD-1.

**S5.**
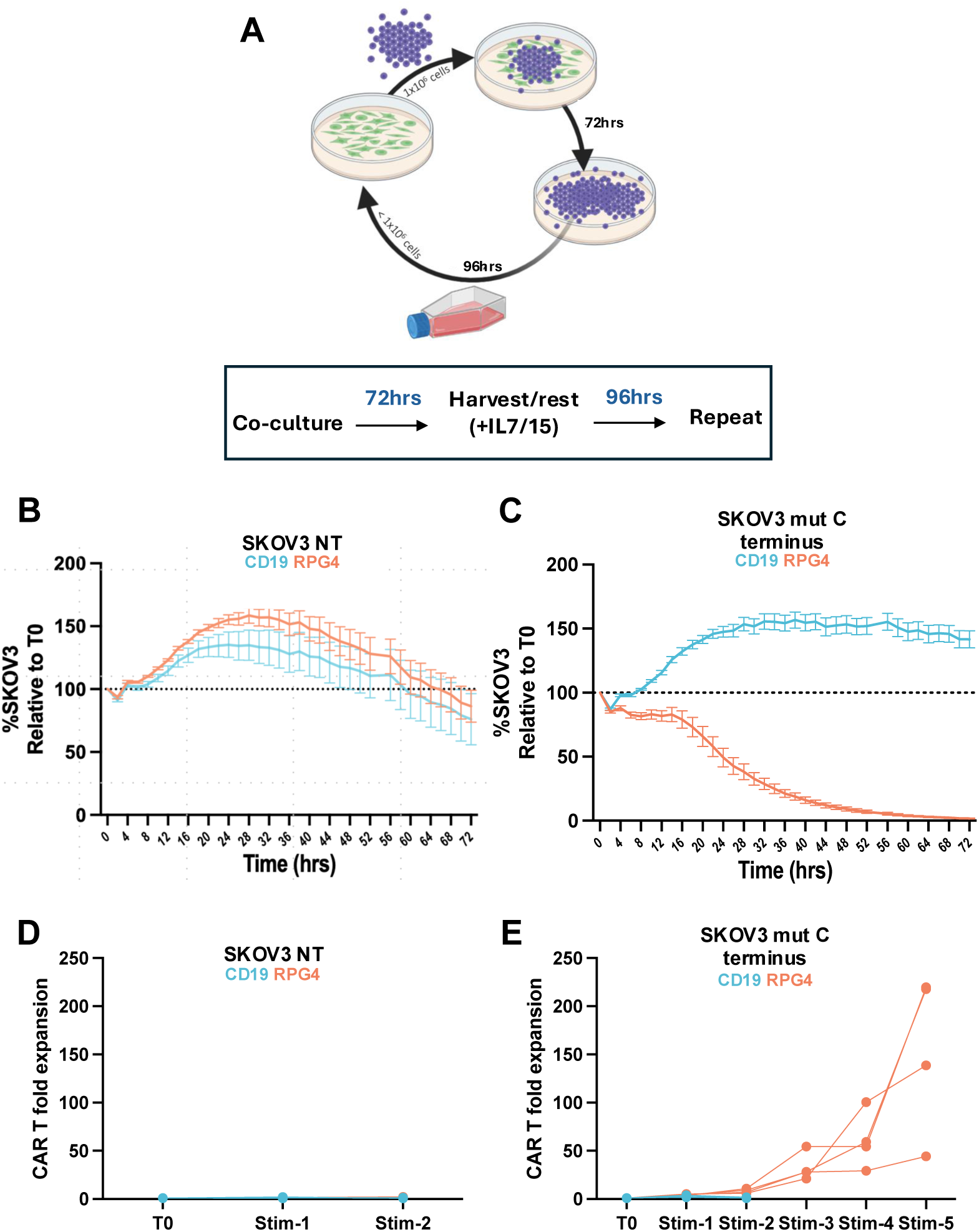
RPG4 CAR T cells show continued efficacy in transient co-stimulation assay and no off-target killing of CALR wildtype, TpoR+ cells in the presence of secreted mutCALR protein. **(A)** Schematic illustrating the experimental setup, showing repeated 72 hour co-culture cycles of CAR T cells (from 4 donors) with target cells, followed by 96 hour rest periods. SKOV3 target cells were plated 24 hours prior to each co-culture, and 1E6 CAR T cells were added at the start of each stimulation. **(b, c)** Example killing curve, from stim-2, depicting % of target cells remaining, relative to time zero, following addition of either RPG4 CAR T cells or CD19 CAR T cells as a negative control. **(d, e)** Theoretical CAR T cell expansion following each 96hr rest period; calculated by multiplying post-restRQR8+ cell counts (10⁶ log) by the initial input for stim-1, or multiplying by the previous fold-expansion for the following stimulations.

**S6.**
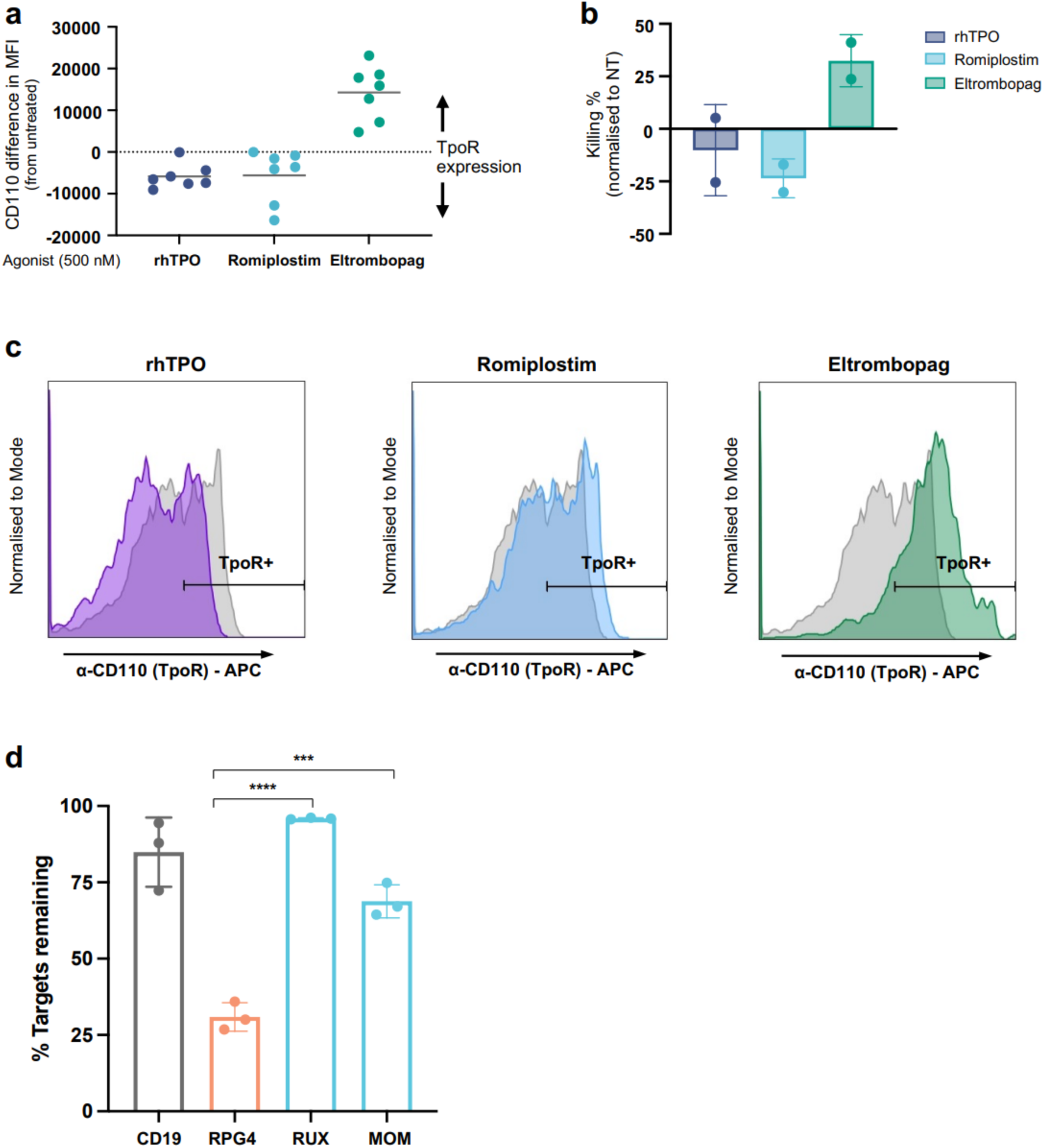
Boosting target expression using TpoR agonists and comparison to JAK inhibitor therapies. **(A)** Change in mean fluorescence intensity (MFI) of CD110 (TpoR) surface expression on Marimo-TpoR cells after overnight treatment with recombinant human TPO (rhTPO), romiplostim, or eltrombopag (500 nM), normalised to untreated control (N=6). **(b)** Change in RPG4 CAR T-mediated killing of Marimo-TpoR cells after overnight pre-treatment with rhTPO, romiplostim, or eltrombopag, normalised to non-transduced T cell condition (NT) (N=2). **(c)** CD110 expression on primary mutCALR+ (type 2) HSPCs following overnight exposure to rhTPO, romiplostim, or eltrombopag (500 nM). (d) Comparative cytotoxicity of RPG4 CAR T to JAK inhibitors ruxolitinib (RUX) 500nM and momelotinib (MOM) 250nM. Chart shows the % of Marimo NT cells remaining (N=3).

**S7.**
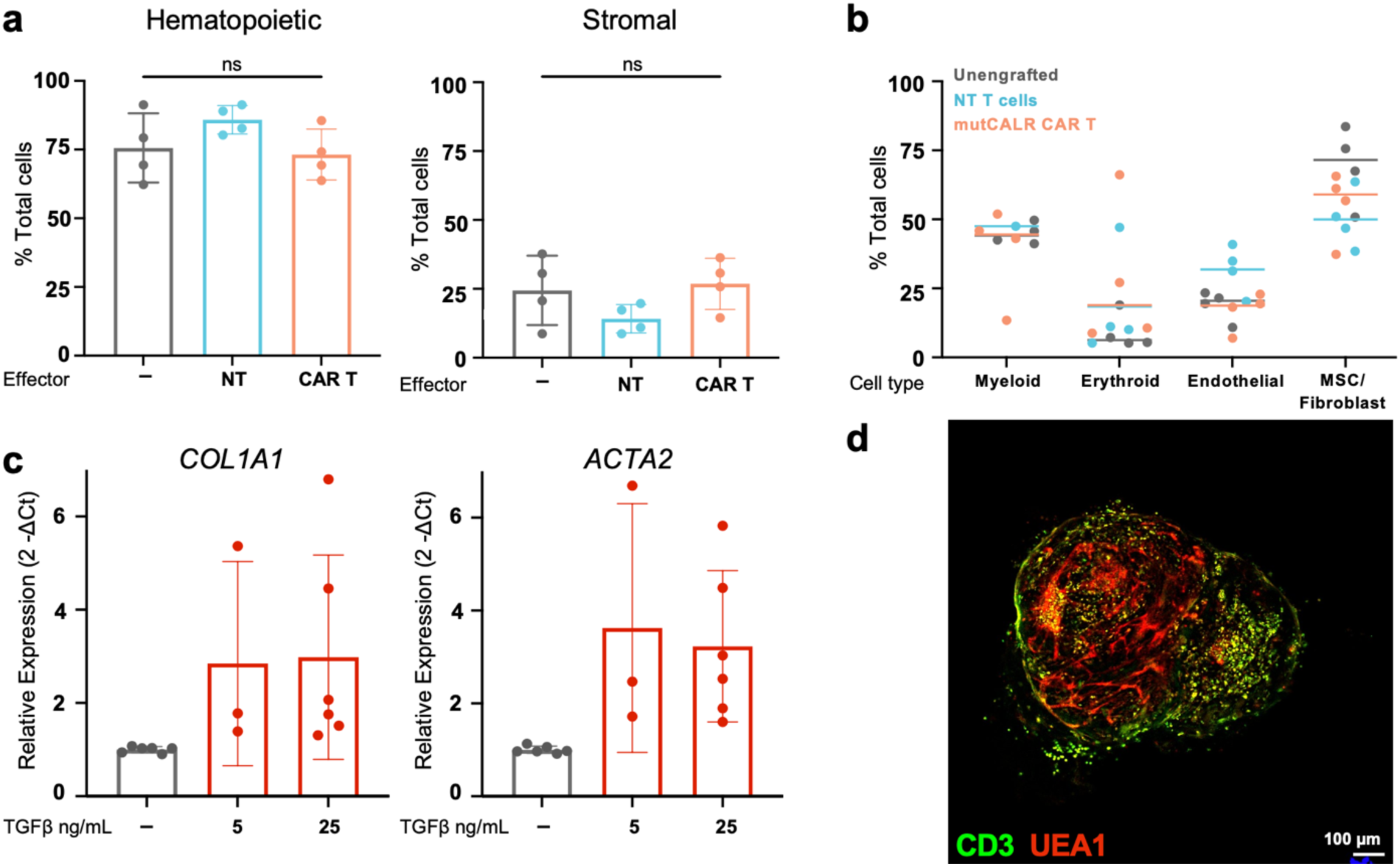
Feasibility of using bone marrow organoids as a platform to assess CAR T cell efficacy in relevant tumor microenvironments. **(a, b)** Characterisation of organoids (grey) after engraftment with non-transduced (NT; blue) T cells or mutCALR-targeting RPG4 CAR T cells (orange). No significant difference in the **(a)** relative proportion of hematopoietic and stromal cells or in **(b)** cell subtypes observed by flow cytometry (Suppl. Table S2). **(c)** Quantification of *COL1A1* (collagen 1) and *ACTA2* (alpha-smooth muscle actin) gene expression in organoids after TGFβ treatment (5 or 25 ng/mL). Each dot represents n = 8-12 organoids, for 3-6 independent experiments. **(d)** Representative image of fibrotic organoid engrafted with CAR T cells, confirming engraftment and dissemination of CAR T cells in a fibrotic marrow model. Green – CD3 (CAR T cells); red – UEA1 (vessels).

**S8.**
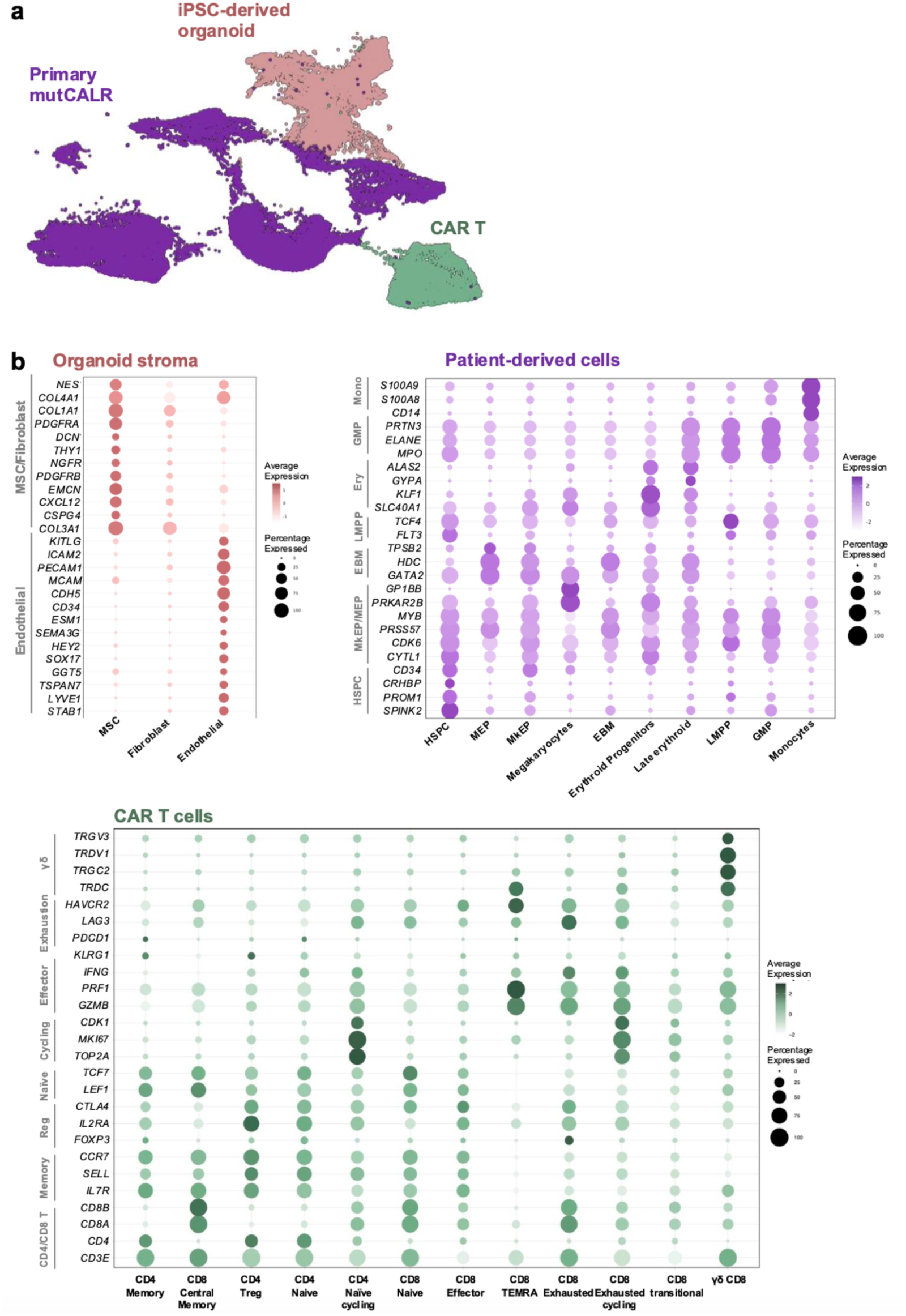
Annotation by genetic de-multiplexing and canonical markers. **(A)** Uniform manifold approximation and projection (UMAP) of genetically de-multiplexed cells, coloured by donor identity (iPSC-derived organoid, pink; primary mutCALR+ myelofibrosis, purple; healthy donor RPG4 CAR T, green). **(b)** Expression of canonical marker genes across cell types from organoid stroma, primary mutCALR+ myelofibrosis, and CAR T cells. Dot size corresponds to percentage of cells expressing each marker and colour intensity indicates average expression level (log-normalised and scaled).

**S9.**
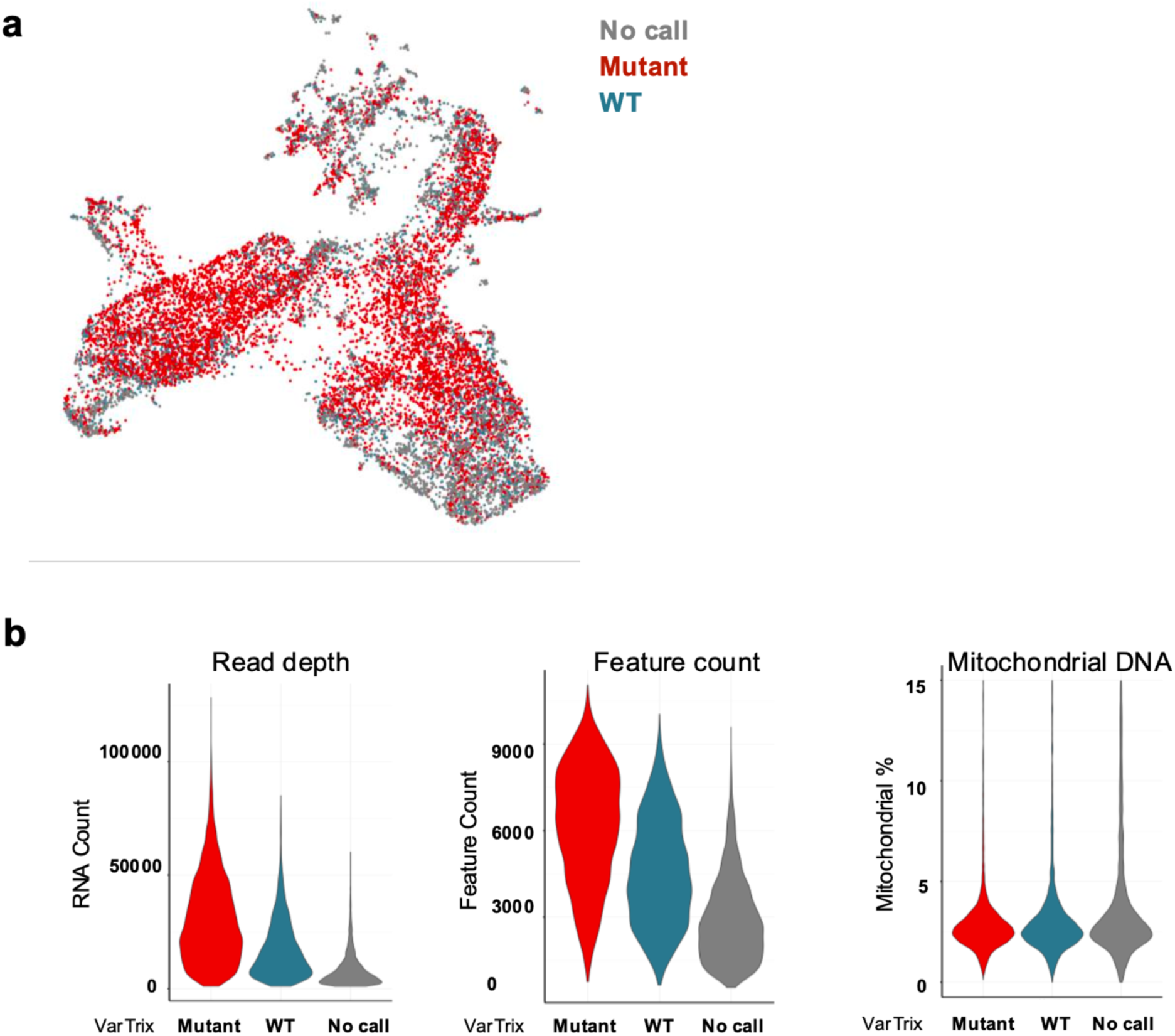
*CALR* mutational analysis of primary mutCALR+ myelofibrosis cells. **(A)** Uniform manifold approximation and projection (UMAP), coloured by type 2 *CALR* mutational status (as determined by Vartrix genotyping analysis). **(b)** Quality metrics of cells, including read depth, number of genes, and percentage of mitochondrial transcripts, categorized by type 2 *CALR* mutational status (mutant, red; wild-type, blue; no call, grey).

**S10.**
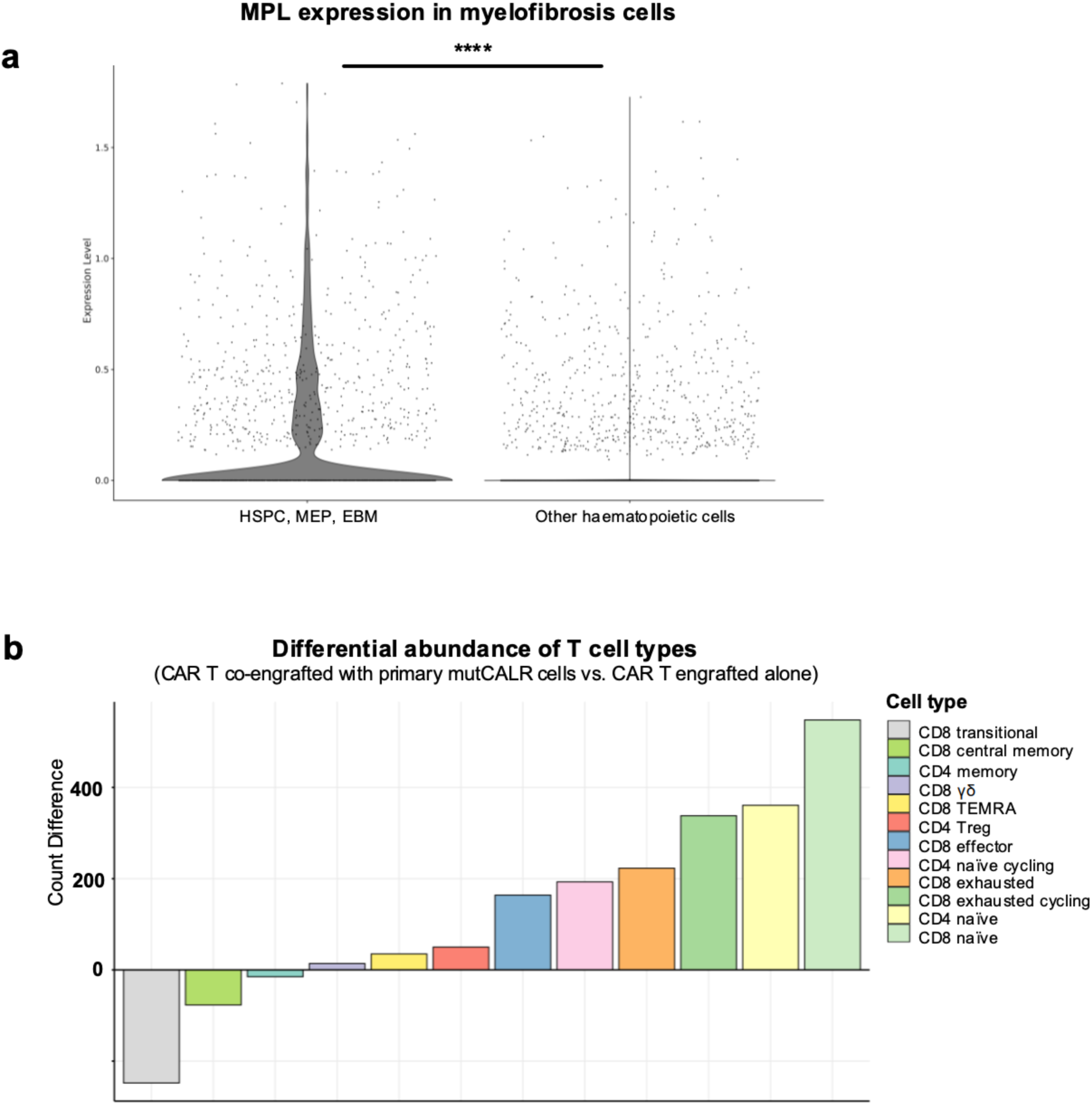
Addition of CAR T cells to chimeroids induces apoptosis of mutCALR myelofibrosis cells that express MPL and leads to expansion of naïve CD4/8+ T cell subsets. **(A)** Expression of MPL for mutCALR+ myelofibrosis cells that were engrafted alone or with RPG4 CAR T cells, with cells grouped by targeted (HSPC, MEP, EBM, left) and non-targeted (all other myelofibrosis cell subsets, right) cell subsets. **** p < 0.0001 for Wilcoxon rank test **(b)** Relative change in T cell phenotypes observed in RPG4 CAR T cells co-engrafted with primary mutCALR+ stem/progenitor cells.

**Table S1.** Clinical characteristics of donors.

| Code number | Date of sample | Diagnosis | Driver mutation | VAF | Secondary mutations | Karyotype | Treatment | Spleen size (cm) | MIPSS score | Hb | WCC | PLTs | % PB blasts |
| --- | --- | --- | --- | --- | --- | --- | --- | --- | --- | --- | --- | --- | --- |
| **Calreticulin mutated** |  |  |  |  |  |  |  |  |  |  |  |  |  |
| 001/115 | 24/07/2019 | ET/Pre-PMF | CALR type 2 | Not reported | Nil | Normal | pre-Peginterferon initiation (previously on Anagrelide) | 14 | Low | 150 | 9.5 | 1223 | 0 |
| 001/128 | 03/10/2022 | PMF | CALR type 1-like (46bp deletion) | 42% | CBL, DNMT3A, GATA (VUS) | 46,XY,-13,+mar[2]/46,idem, add(3)(q27?), del(20)(q11.2q13.3)[7]/45,idem, add(3)(q27?), del(20)(q11.2q13.3),-22[3] | Ruxolitinib 10mg bd plus CPI-0610 75mg | Splenectomy | High | 80 | 1.48 | 49 | 19 |
| 001/178 | 14/01/2021 | PMF | CALR type 2 | 32% | Nil | Normal | Nil | 15 | Low | 126 | 7.63 | 958 | 0 |
| 001/182 | 13/09/2022 | PMF | CALR type 1 | 24% | CHEK2 (VUS) | Normal | Peginterferon | 13 | Low | 141 | 4.18 | 424 | 0 |
| 001/236 | 21/11/2022 | PMF | CALR type 2 | 50% | ASXL1, EZH2 (80%) | Normal | Nil | 26 | Intermediate | 110 | 17.4 | 780 | 3 |
| 001/270 | 01/04/2022 | PMF | CALR type 1 | 41% | ASXL1, BRAF, GNAS, TET2, ZRSR2 | Not known | Ruxolitinib (Freedom2) | 16 | High | 88 | 3.35 | 84 | 1 |
| 001/374N | 14/04/2024 | Post ET MF | CALR type 2 | 44% | ASXL1, ETV6, ZRSR2 | Failed | Nil | 17.3 | Intermediate | 76 | 13.68 | 825 | 0 |
| 001/391 | 12/05/2024 | ET | CALR type 2 | 25% | PPM1D | Not available | Hydroxycarbamide | 12 | Low | 135 | 3.55 | 395 | 0 |
| 001/366 | 06/02/2024 | PMF | CALR type 2 like | 39% | NFE2 VUS | Normal | Nil | 13.9 | Intermediate | 114 | 10.28 | 172 | 0 |
| 001/392 MFb1 (IHC) | 03/06/2024 | PMF | CALR type 1 | 50% | CSF3R, PPM1D | Normal | Hydroxycarbamide | 12 | Low | 113 | 3.02 | 347 | 0 |
| 001/098 | 18/07/2023 | PMF | CALR type 2 | 58% | ASXL1 | Not known | Ruxolitinib | 16.4 | Intermediate | 98 | 9.55 | 364 | 13 |
| 001/351 | 12/06/2023 | Post ET MF | CALR type 1 | 46% | ASXL1 | Normal | Hydroxycarbamide | 14 | High | 119 | 28.45 | 313 | 9 |
| ETb (IHC) | 15/05/2023 | ET | CALR type 2 | Not reported | Nil | Not available | Hydroxycarbamide | 12 | N/A | 105 | 4.38 | 548 | 0 |
| 003/035 | 29/08/2018 | BP-MPN | CALR | 26% | EZH2, NRAS | CNN-LOH on 7q | Azacitidine/Ruxolitinib (PHAZAR) | N/A | N/A | 103 | 7 | 378 | 22 |
| **JAK2 mutated** |  |  |  |  |  |  |  |  |  |  |  |  |  |
| 001/283 | 22/08/2023 | PMF | JAK2 V617F | >90% | ASXL1 x2 | Not available | Mometotinib | Not available | Int-2 | 80 | 61 | 101 | 3 |
| 001/204N | 21/02/2025 | Post PV MF | JAK2 V617F | 95% | Nil | Not available | Hydroxycarbamide Navitoclax | Splenectomy | Int-1 | 129 | 54 | 1019 | 0 |
| 001/159 | 20/05/2020 | Pre-PMF/PMF | JAK2 V617F | 40% | ASXL1, EZH2, ZRSR1 | Not available | Nil | Not available | Int-1 | 111 | 7.17 | 600 | 0 |
| 001/177 | 23/03/2023 | PMF | JAK2 V617F | 70% | ASXL1, SRSF2 TET2, TET2 (VUS) | Not available | Ruxolitinib plus Navitoclax/Placebo (TRANSFORM-1) | Not available | Int-2 | 118 | 8.19 | 129 | 0 |
| 001/365 | 27/02/2025 | PV | JAK2 V617F | 20% | BCOR | Not available | Nil | Not available | N/A | 150 | 8.22 | 709 | 0 |
| 001/395 | 29/02/2024 | PMF | JAK2 V617F | Not reported | ASXL1, ETV6, PHF6, SETBP1 | Not available | Mometotinib | Not available | Int-2 | 91 | 62.7 | 186 | 8 |
| MFaJAK (IHC) | 22/02/2024 | Post PV MF | JAK2 V617F | Not reported | U2AF1 | del5 | Hydroxycarbamide | 19.4 | Int-2 | 70 | 26.7 | 22 | 2 |
| MFbJAK (IHC) | 09/01/2024 | PMF | JAK2 V617F | 70% | Nil | Normal | Nil | 19 | Int-1 | 109 | 19.51 | 257 | 0 |

**Table S2.** Antibodies for flow cytometry and imaging.

| Flow Cytometry Antibodies |  |  |  |  |
| --- | --- | --- | --- | --- |
| Antigen | Fluorophore | Supplier | CAT# | Dilution |
| hCCR7 | FITC | BioLegend | 353216 | 1:100 |
| hCD25 | APC/Cy7 | BioLegend | 302614 | 1:100 |
| hCD3 | PE/Cy7 | BioLegend | 981010 | 1:100 |
| hCD34 | FITC | R&D Systems | FAB7227G | 1:100 |
| hCD34 | APC | BioLegend | 378606 | 1:100 |
| hCD45RA | AF700 | BioLegend | 304120 | 1:100 |
| hCD62L | PE | BioLegend | 304805 | 1:100 |
| hCD69 | APC/Cy7 | BioLegend | 310914 | 1:100 |
| hCD95 | Pacific Blue | BioLegend | 305619 | 1:100 |
| hLag3 | PerCP/Cy5.5 | BioLegend | 369312 | 1:100 |
| hPD1 | PE | BioLegend | 379210 | 1:100 |
| hTIM3 | APC | BioLegend | 345012 | 1:100 |
| mlgG2a | PE | Jackson Laboratories | 115-115-206 | 1:1000 |
| Viability dye | ef780 | Thermofisher | 65-0865-14 | 1:1000 |
| Viability dye | SYTOX™ Blue | Invitrogen | S34857 | 1:1000 |
| Viability dye | 7AAD | BioLegend | 420404 | 3:100 |

| Memory panel |  |  |  |  |
| --- | --- | --- | --- | --- |
| hCCR7 | FITC | BioLegend | 353216 | 1:100 |
| hCD3 | PE/Cy7 | BioLegend | 981010 | 1:100 |
| hCD34 | APC | BioLegend | 378606 | 1:100 |
| hCD45RA | AF700 | BioLegend | 304120 | 1:100 |
| hCD62L | PE | BioLegend | 304805 | 1:100 |
| hCD95 | Pacific Blue | BioLegend | 305619 | 1:100 |
| Viability dye | 7AAD | BioLegend | 420404 | 3:100 |

| Exhaustion panel |  |  |  |  |
| --- | --- | --- | --- | --- |
| hCD3 | PE/Cy7 | BioLegend | 981010 | 1:100 |
| hCD34 | FITC | R&D Systems | FAB7227G | 1:100 |
| hLag3 | PerCP/Cy5.5 | BioLegend | 369312 | 1:100 |
| hPD1 | PE | BioLegend | 379210 | 1:100 |
| hTIM3 | APC | BioLegend | 345012 | 1:100 |
| Viability dye | 7AAD | BioLegend | 420404 | 3:100 |

| Killing assay panel vs cell lines |  |  |  |  |
| --- | --- | --- | --- | --- |
| hCD3 | PE/Cy7 | BioLegend | 981010 | 1:100 |
| hCD34 | APC | BioLegend | 378606 | 1:100 |
| Viability dye | ef780 | Thermofisher | 65-0865-14 | 1:1000 |
| CALR targets | eGFP |  |  |  |
| TpoR targets | eBFP |  |  |  |

| Activation panel |  |  |  |  |
| --- | --- | --- | --- | --- |
| hCD3 | PE/Cy7 | BioLegend | 981010 | 1:100 |
| hCD34 | APC | BioLegend | 378606 | 1:100 |
| Viability dye | SYTOX™ Blue | Invitrogen | S34857 | 1:1000 |
| hCD69 (24hrs) | APC/Cy7 | BioLegend | 310914 | 1:100 |
| hCD25 (72hrs) | APC/Cy7 | BioLegend | 302614 | 1:100 |

| Organoid immunophenotyping panel |  |  |  |  |
| --- | --- | --- | --- | --- |
| hCD71 | BV650 | BioLegend | 334116 | 1:100 |
| hCD235 | PE/Cy7 | BioLegend | 306620 | 1:400 |
| hCD11b | Alexa 700 | BioLegend | 301355 | 1:200 |
| hCD14 | Alexa 700 | BioLegend | 367113 | 1:200 |
| hCD45 | FITC/AF488 | BioLegend | 304017 | 1:100 |
| hCD90 | PE/Cy5 | BioLegend | 328112 | 1:200 |
| hCD31 | APCCy7 | BioLegend | 303120 | 1:100 |
| Viability dye | DAPI | Life Technologies | D3571 | 1:50k |

| TpoR surface expression |  |  |  |  |
| --- | --- | --- | --- | --- |
| hCD110 | APC | Miltenyi Biotec | 130-128-223 | 1:100 |
| IgG1 Isotype control | APC | BioLegend | 403506 | 1:400 |

| Imaging antibodies |  |  |  |  |
| --- | --- | --- | --- | --- |
| Primary | Host/Isotype/Class | Supplier | CAT# | Dilution |
| hCD3 | Mouse/IgG1, κ/monoclonal | Invitrogen | MA5-12577 | 2:100 |
| UEA1 | Biotinylated | Vector Labs | B-1065 | 5:100 |
| Secondary | Fluorophore | Supplier | CAT# | Dilution |
| Goat a-Mouse IgG2a | Alexa Fluor 488 | ThermoFisher | A-21131 | 1:250 |
| Streptavidin | Alexa Fluor 568 | Invitrogen | S11226 | 1:250 |

